# Distinctive chaperonopathy in skeletal muscle associated with the dominant variant in *DNAJB4*

**DOI:** 10.1101/2022.07.26.501446

**Authors:** Michio Inoue, Satoru Noguchi, Yukiko U. Inoue, Aritoshi Iida, Megumu Ogawa, Rocio Bengoechea, Sara K. Pittman, Shinichiro Hayashi, Kazuki Watanabe, Yasushi Hosoi, Terunori Sano, Masaki Takao, Yasushi Oya, Yuji Takahashi, Hiroaki Miyajima, Conrad C. Weihl, Takayoshi Inoue, Ichizo Nishino

**Affiliations:** Department of Neuromuscular Research, National Institute of Neuroscience, National Center of Neurology and Psychiatry, Tokyo, Japan; Medical Genome Center, National Center of Neurology and Psychiatry, Tokyo, Japan; Department of Neurology, Washington University School of Medicine, Saint Louis, USA; Department of Biochemistry and Cellular Biology, National Institute of Neuroscience, National Center of Neurology and Psychiatry, Tokyo, Japan; First Department of Medicine / Department of Neurology, Hamamatsu University School of Medicine, Hamamatsu, Japan; Department of Laboratory Medicine, National Center Hospital, National Center of Neurology and Psychiatry, Japan; Department of Neurology, National Center Hospital, National Center of Neurology and Psychiatry, Tokyo, Japan

## Abstract

DnaJ homolog, subfamily B, member 4, a member of the heat shock protein 40 chaperones encoded by *DNAJB4*, is highly expressed in myofibers. We identified a heterozygous c.270 T>A (p.F90L) variant in *DNAJB4* in a family with a dominantly inherited distal myopathy, in which affected members have specific features on muscle pathology represented by the presence of cytoplasmic inclusions and the accumulation of desmin, p62, HSP70 and DNAJB4 predominantly in type 1 fibers. Both Dnajb4- F90L knock-in and knockout mice developed muscle weakness and recapitulated the patient muscle pathology in the soleus muscle, where DNAJB4 has the highest expression. These data indicate that the identified variant is causative resulting in defective chaperone function and selective muscle degeneration in specific muscle fibers. This study demonstrates the importance of DNAJB4 in skeletal muscle proteostasis by identifying the associated chaperonopathy.

## Introduction

Molecular chaperones are proteins that assist in the conformational refolding or degradation of misfolded proteins, thus they determine the fate of client proteins and contribute to the maintenance of cellular protein homeostasis (known as proteostasis)^1, 2^. In skeletal muscle, chaperones are highly expressed and work to protect cells from several stresses^3^. The small heat shock proteins and nucleotide exchange factors, or co- chaperones, work together with molecular chaperones on client protein recognition and the regulation of ATPase activity^4, 5^.

Mutations in chaperone or co-chaperone genes cause neuromuscular diseases, termed chaperonopathies^1, 6^. Previous studies found that mutations in *CRYAB*^7^, *HSPB8*^8^, *BAG3*^9^*, DNAJB6*^10^, or *SIL1*^11, 12^ cause protein aggregate myopathies including myofibrillar myopathy, rimmed vacuolar myopathy, limb girdle muscular dystrophy (LGMD) D1 or Marinesco-Sjögren syndrome, respectively. A common pathogenic process is the functional disturbance of the molecular chaperone systems by the genetic variants leading to aggregation of misfolded proteins in myofibers and subsequent muscle degeneration^6, 13^. However, the precise pathomechanism, including the identity of client proteins, the contents and properties of the cytoplasmic aggregates, their toxicities and consequent effects, and the response of myofibers, has not been completely clarified in these myopathies.

On muscle pathology, the diseases representing the presence of protein aggregation with myofibril degeneration in myofibers are collectively referred to as protein aggregate myopathies or myofibrillar myopathies^14^. Recent widespread use of next generation sequencing has enabled the identification of causative genes in protein aggregate myopathies^6, 15^. However, many patients remain to be genetically diagnosed.

For example, 13 causative genes are known in protein aggregate/myofibrillar myopathy to date^16^, yet the diagnosed rate of patients with this disease is only 30%^17^.

DNAJB4, a class II J-domain protein, or HSP40, is a co-chaperone of the ubiquitous Hsp70 chaperone and is highly expressed in striated muscles. Its homologous protein, DNAJB6, is also expressed in striated muscles and dominant DNAJB6 mutations cause LGMDD1, in which muscles have protein aggregate and vacuolar pathology^10^. To date, no human disease related to *DNAJB4* has been reported. In this study, we identified a pathogenic variant in *DNAJB4* in patients with clinically, a distal myopathy and myopathologically, a protein aggregate myopathy. Moreover, we established the DNAJB4 variant as pathogenic using cellular and animal models.

Furthermore, we generated two animal models, *Dnajb4* variant-knock-in (KI) and knockout (KO) mice, which recapitulated the phenotypes of the patients, suggesting that the dysproteostasis is the basic mechanism of this disease.

## Results

### Clinicopathological features of the patients

The proband (II-5) (Fig. 1a) was the third-born child from non-consanguineous parents. Her father, two brothers, sister, and the son of the second brother had similar symptoms, suggesting an autosomal dominant mode of inheritance. In affected individuals, motor development in childhood was unremarkable. In the third to fifth decade of life, they presented with asymmetric thumb and grip weakness resulting in atrophy of thenar and hypothenar muscles (Fig. 1b) and finger contractures. Subsequent progression included inability to stand on tiptoes and the weakness and atrophy of distal leg muscles. Further, they ultimately developed asymmetric involvement of proximal muscles, respiratory insufficiency, scoliosis, and loss of ambulation in the seventh decade (Supplementary Table 1).

**Fig. 1.**
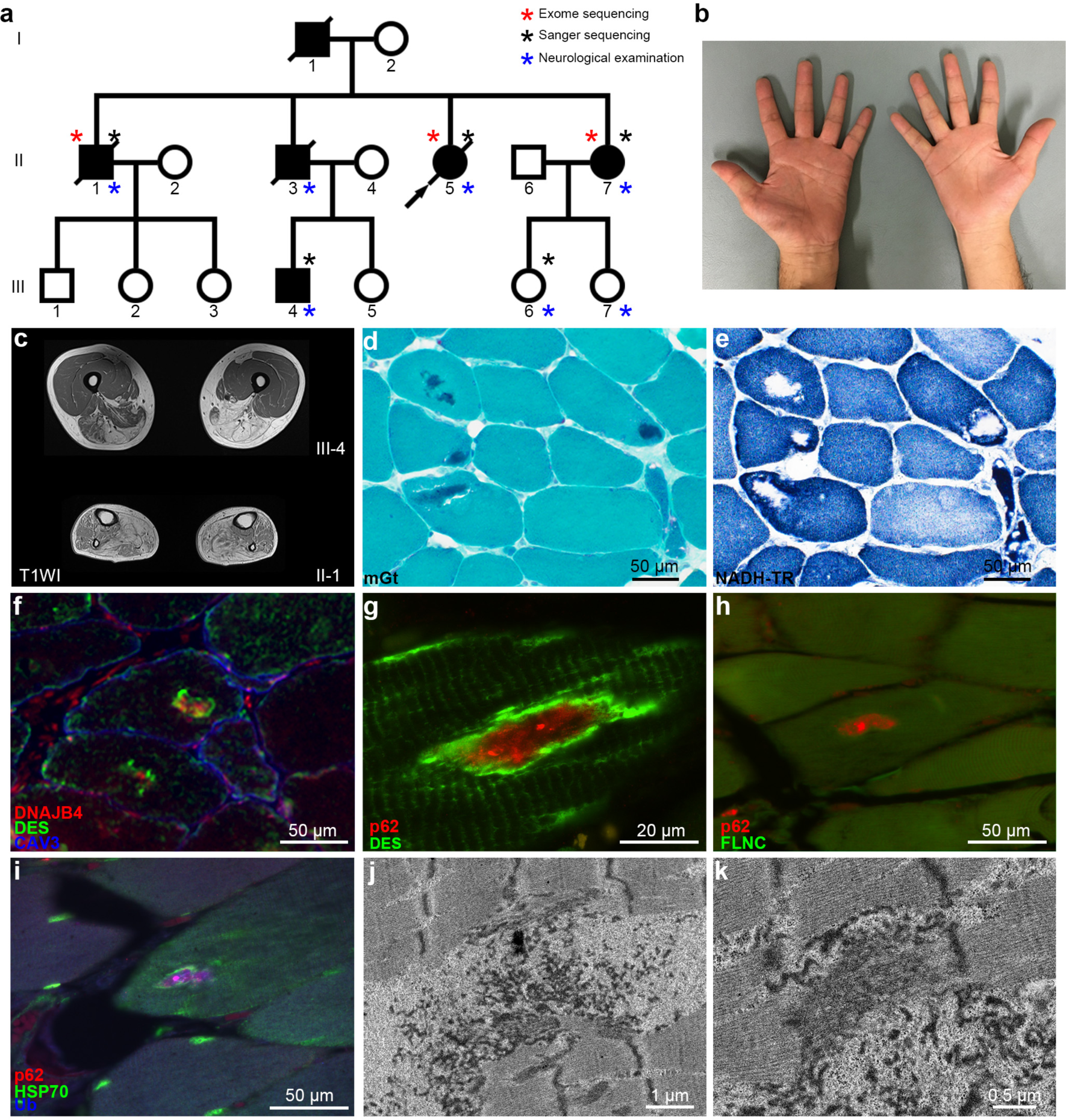
Clinical, myopathological, and ultrastructural findings from a family with dominantly inherited myopathy. **a** A pedigree showing dominantly inherited myopathy. An arrow denotes the proband and symbols with a slash denote deceased members of the family. **b** The picture of hands (III-4) at 49 years of age showing asymmetric atrophy of thenar and hypothenar muscles. **c** Magnetic resonance imaging of thigh muscles of III-4 at 49 years of age and lower legs of II-1 at 71 years of age (T1-weighted image, T1WI). **d-k** Muscle pathology, immunohistochemistry, and electron microscopic pictures of muscles from II-5. **d** Modified Gomori trichrome (mGt). **e** Nicotinamide adenine dinucleotide dehydrogenase-tetrazolium reductase (NADH- TR) staining is serial section of **d**. **f**-**i** Immunofluorescence staining for the indicated antibodies. Muscle membranes are stained by anti-caveolin-3 (CAV3) antibodies (blue). **j** Electron microscopic photograph of muscle from II-5, showing electron dense inclusion. **k** High magnification image showing degenerating thin and thick filaments. Scale bar = 50 μm (**d**-**f**, **h**, **i**), 20 μm (**g**), 1 μm (**j**), 0.5 μm (**k**). DES, desmin; FLNC, filamin C; HSP70, heat shock protein 70; NADH-TR, nicotinamide adenine dinucleotide dehydrogenase-tetrazolium reductase; mGt, modified Gomori trichrome; Ub, ubiquitin

Serum creatine kinase levels were elevated (646 – 1285 U/mL; normal levels, 42 – 164 U/mL). Muscle MRI and CT from three patients showed asymmetric muscle involvement, as evidenced by various fat replacement in the adductor magnus, adductor longus, biceps femoris, rectus femoris, and gastrocnemius. This contrasts symmetric involvement in the semitendinosus, semimembranosus, gracilis, soleus, and paraspinal muscles (Fig. 1c and Supplementary Fig. 1).

Muscle pathology showed moderate variation in fiber size, occasional rimmed vacuoles in atrophic fibers, and scattered fibers with cytoplasmic inclusions on modified Gomori trichrome (mGt) (Fig. 1d). On serial sections, nicotinamide adenine dinucleotide dehydrogenase-tetrazolium reductase (NADH-TR) revealed that scattered fibers with cores corresponded to the cytoplasmic inclusions that found predominantly in dark stained fibers consistent with type 1 fibers (Fig. 1e).

### Identification of a DNAJB4 missense variant and accumulation of DNAJB4 in patient muscle

We performed whole exome sequencing on the proband and two siblings (Fig, 1a II-1, II-5, II-7). We focused on the 17 variants shared by the three affected family members (Supplementary Table 2). Among the 17 filtered variants, we evaluated cosegregation among affected (II-5, II-7, III-4) and unaffected individuals (III-6) by Sanger sequencing. We identified one heterozygous nonsynonymous variant (c.270T>A, p.F90L) in *DNAJB4* (NM_007034) and one heterozygous synonymous variant (c.1612C>T, p.L538L) in *DDX58* (NM_014314). No variants in other genes associated with muscle diseases were found. We further analyzed the missense variant (c.270T>A) in *DNAJB4*. The phenylalanine at 90 in the glycine/phenylalanine-rich domain is not only highly conserved but also corresponds to the F93 residue in *DNAJB6*, which when mutated to leucine or isoleucine causes LGMDD1^10^ (Fig 2a, b, c).

**Fig. 2.**
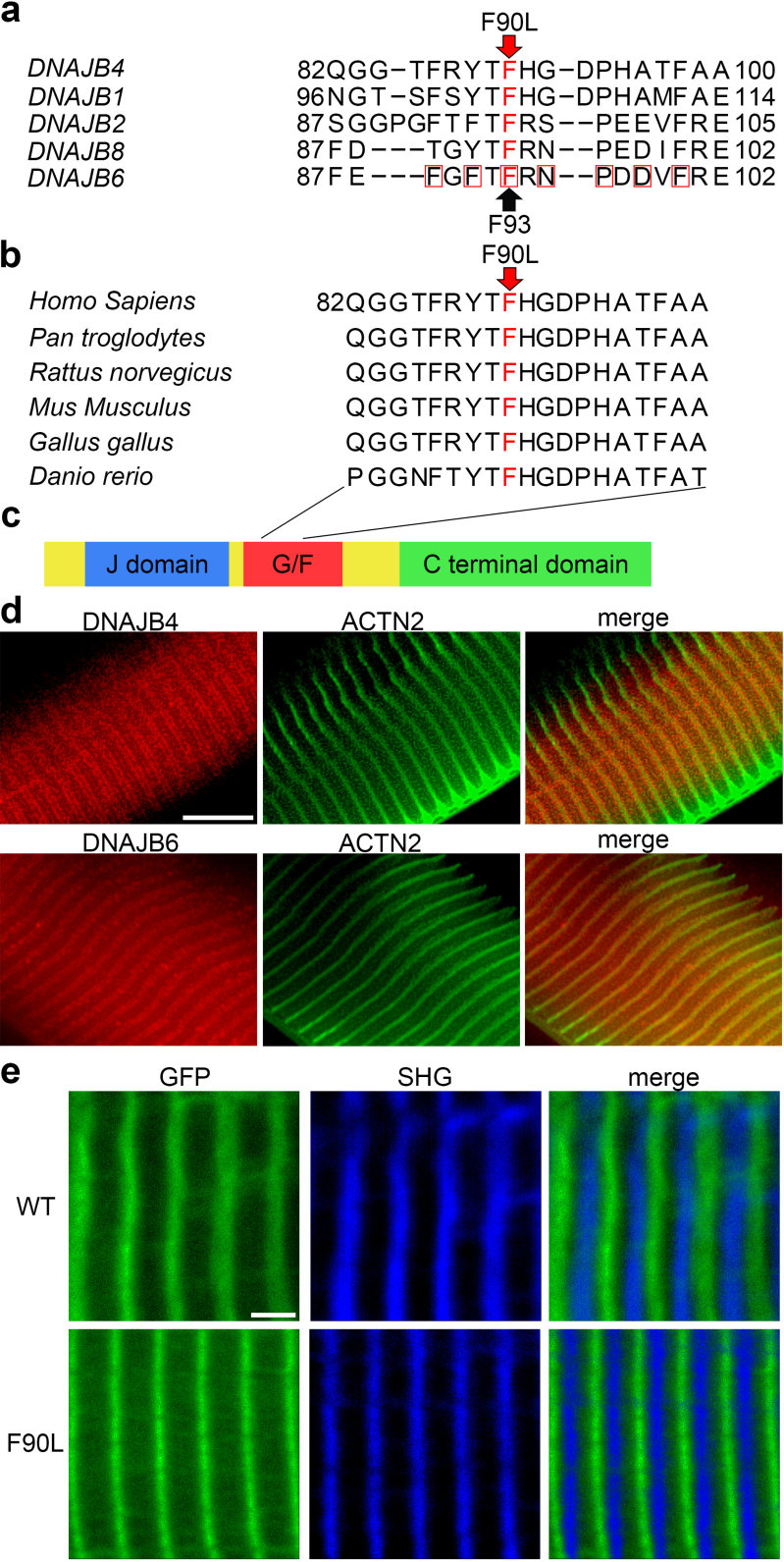
High homology of the affected residue in the G/F domain of DNAJB4 and localization of DNAJB4. **a** Sequence alignment of human DNAJB4 with paralogs performed using Protein BLAST showing the disease-affected residue in the human myopathy patients is highly conserved. Red squares denote the residues in *DNAJB6*, which when mutated cause myopathy. **b** Sequence alignment of DNAJB4 orthologues showing the affected phenylalanine residue is evolutionarily conserved. **c** Schema of the domain structure of DNAJB proteins. **d** Localization of DNAJB4 or DNAJB6 in single fiber of the soleus muscles of wild-type C57BL/6J mice. Both DNAJB4 and DNAJB6 are colocalized with alpha-actinin-2 (ACTN2) or the Z-discs. **e** GFP tagged wild-type DNAJB4 (WT) or DNAJB4-90L (F90L) were electroporated in the mouse the flexor digitorum brevis muscles and mouse foot pad was imaged via two-photon microscopy. Second harmonic generation (SHG) was used to visualize the A-band for a reference point. Scale bar = 10 μm (**d**), 2 μm (**e**)

DNAJB4 inclusions were surrounded by desmin in the center of myofibers (Fig. 1f). High magnification of longitudinal sections along muscle fibers revealed desmin similarly accumulated surrounding p62 aggregates. These aggregates were quite large spanning more than 30 sarcomeres (Fig. 1g). Filamin C and HSP70 were also colocalized with p62-positive deposits (Fig. 1h, i). Ultrastructural analysis of the patient muscle showed woolly inclusions whose electron density was similar or lower compared to the adjacent disorganized Z-discs (Fig. 1j). Magnified image showed appearing-normal Z-disc adjacent to degenerated thin and thick filaments (arrow), suggesting degeneration may start from myofibrils (Fig. 1k).

To investigate the localization of DNAJB4 in normal myofibers, we performed single fiber analysis of the soleus muscles from wild-type mice, and found that DNAJB4 was localized to the vicinity of the Z-disc similar to DNAJB6, where ACTN2 was colocalized (Fig 2d). Furthermore, to analyze whether the F90L mutation induces a change in protein localization, we performed second harmonic generation imaging using the flexor digitorum brevis (FDB) muscles electroporated with GFP-tagged wild-type or DNAJB4-F90L, and found that the F90L variant in *DNAJB4* did not alter its localization on the Z-disc (Fig. 2e).

### Dnajb4^-/-^ mice developed muscle weakness

We generated *Dnajb4* KI mice harboring c.270C>A mutation (Dnajb4^F90L/+^) and KO mice harboring c.269_270insGT mutation (Dnajb4^+/-^), which is predicted to cause a frameshift at 270 and generate a premature stop codon at 384, using CRISPR/cas9 gene editing (Supplementary Fig. 2a). The mouse genotypes were confirmed by Sanger sequencing (Supplementary Fig. 2b). Heterozygous Dnajb4^F90L/+^ mice could be bred to generate homozygous Dnajb4^F90L/F90L^ mice. Heterozygous Dnajb4^+/-^ also enabled the generation of homozygous Dnajb4^-/-^ mice. Dnajb4^-/-^ mice had significantly reduced DNAJB4 protein detected by western blotting of quadriceps muscle lysates (Supplementary Fig. 2c).

Heterozygous Dnajb4^F90L/+^ mice had no difference in body weight up to 13 months compared to the wild-type littermates (Dnajb4^+/+^) (Fig. 3a). Grip strength of Dnajb4^F90L/+^ and Dnajb4^F90L/F90L^ mice did not show a decrease compared to wild type, while Dnajb4^-/-^ mice showed evidence of grip weakness at five months of age (Fig. 3b).

**Fig. 3.**
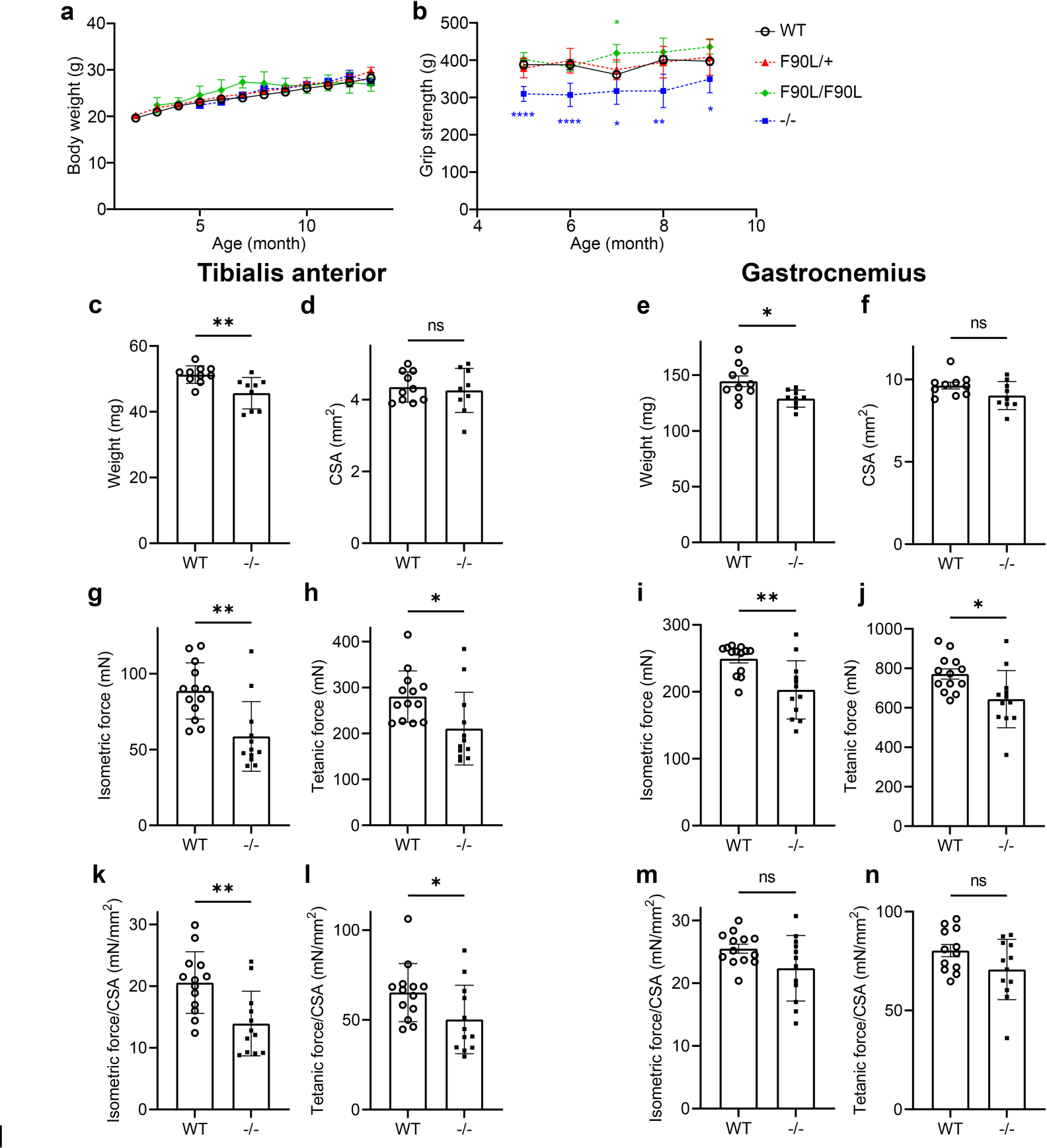
Knockout of Dnajb4 in mouse results in muscle weakness. **a** Average body weight (g) at 2 to 13 months of age in Dnajb4^F90L/+^ (F90L/+) (n = 4-7 per aged group), Dnajb4^F90L/F90L^ (F90L/F90L) (n = 4-11 per aged group), Dnajb4^-/-^ (-/-) (n = 10-23 per aged group), and Dnajb4^+/+^ (WT) mice (n = 5-6 per aged group). **b** Average score for the grip strength at 5 to 9 months in in Dnajb4^F90L/+^ (n = 7-12 per aged group), Dnajb4^F90L/F90L^ (n = 4-11 per aged group), Dnajb4^-/-^ (n = 10-19 per aged group), and Dnajb4^+/+^ mice (n = 5-6 per aged group). Muscle weight (**c**, **e**), CSA (cross sectional area) (**d**, **f**), and contractile forces (**g**-**n**) of tibialis anterior muscles (**c**, **d**, **g**, **h**, **k**, **l**) and gastrocnemius (**e**, **f**, **i**, **j**, **m**, **n**) at 13 months of age (n = 12-13 per each group): (**g**, **i**) isometric twitch force, (**h**, **j**) maximum tetanic force, (**k**, **m**) specific isometric force normalized by CSA, (**l, n**), and specific tetanic force normalized by CSA. Values from each group are expressed as mean ± SEM. A paired Student’s t-test for comparison between two groups or a multiple testing comparison with multiple testing correction (Dunnett) was performed. *p < 0.05, **p < 0.01, ****p < 0.0001, and ns, not significant

We analyzed muscle contraction of the tibialis anterior and gastrocnemius muscles, and found decreased muscle mass, peak isometric force, and maximum tetanic force in Dnajb4^-/-^ mice at the age of 13 months (Fig. 3c-n). In contrast, no change was seen in Dnajb4^F90L/F90L^ mice at 13 months of age (data not shown). These results suggest that DNAJB4 may be essential for the contraction of skeletal muscle.

To explore a later age phenotype, we extended the scope of our analysis to 28 months. Dnajb4^F90L/+^ or Dnajb4^-/-^ mice showed no change in life span (Fig. 4a), body weight (Fig. 4b), or muscle contraction of the tibialis anterior and gastrocnemius muscles (Supplementary Fig. 3) compared to Dnajb4^+/+^. In contrast, Dnajb4^F90L/+^ mice had a mild decrease in grip strength at 28 months of age (Fig. 4c).

**Fig. 4.**
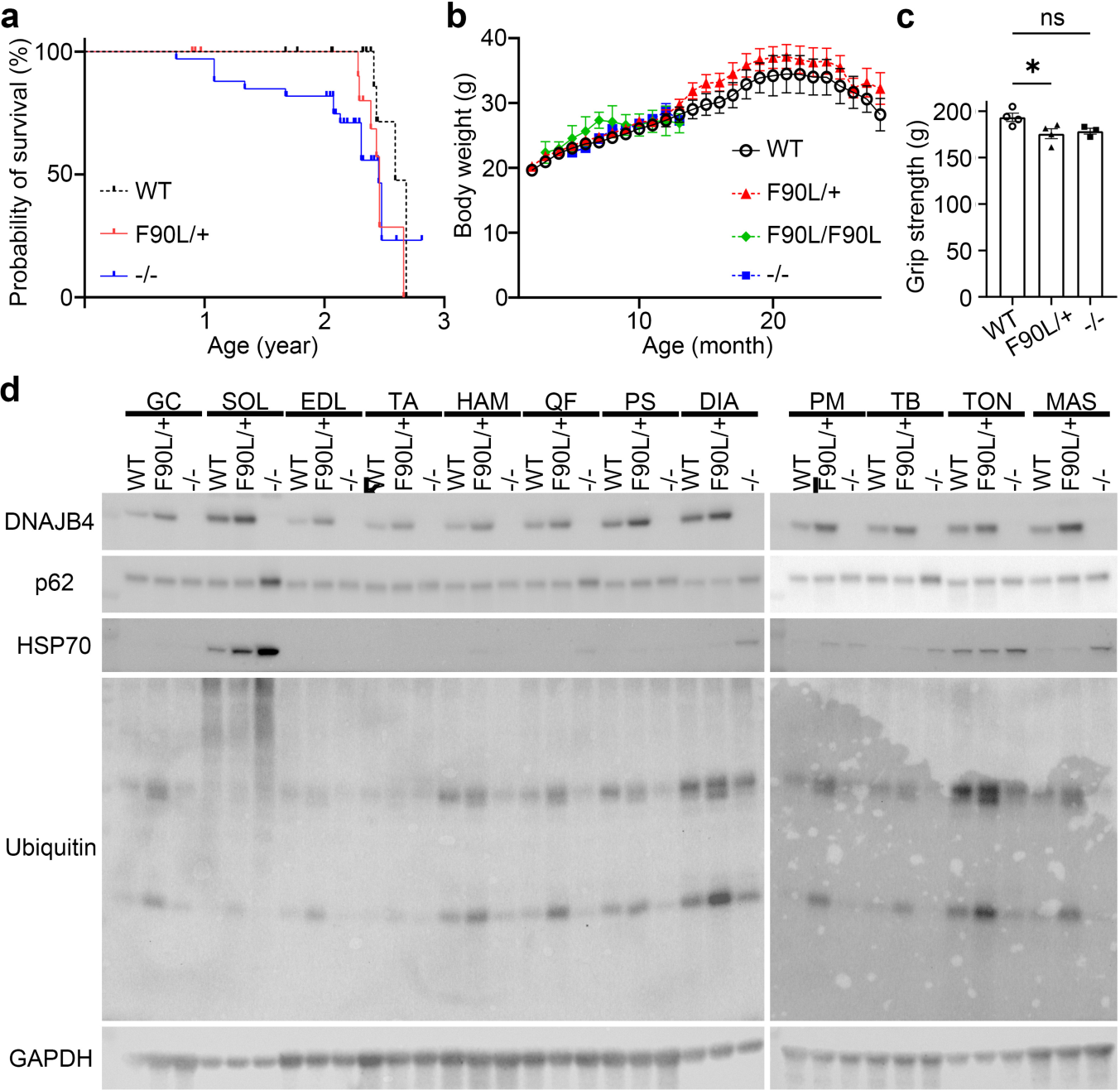
Physiological effect by and muscle susceptibility to Dnajb4 dysfunction in mice. **a** Survival rate of Dnajb4^F90L/+^ (F90L/+) (n = 14), Dnajb4^-/-^ (-/-) (n = 33), and Dnajb4^+/+^ (WT) mice (n = 16) shows no significant difference between groups (F90L vs WT, p = 0.10; -/- vs WT, p = 0.086). Tick marks indicate censored data. P value was obtained from the log rank test. **b** Average body weight at 2 to 28 months of age in Dnajb4^F90L/+^ (n = 4-7 per aged group), Dnajb4^F90L/F90L^ (F90L/F90L) (n = 4-11 per aged group), Dnajb4^-/-^ (n = 10-23 per aged group), and Dnajb4^+/+^ mice (n = 5-6 per aged group). **c** Average score for the grip strength at 28 months in in Dnajb4^F90L/+^ (n = 4), Dnajb4^-/-^ (n = 3), and Dnajb4^+/+^ mice (n = 4). A multiple testing comparison with multiple testing correction (Dunnett) was performed. *p < 0.05 and ns, not significant. **d** Immunoblot of lysates from indicated muscles of Dnajb4^+/+^, Dnajb4^F90L/+^, and Dnajb4^-/-^ mice at 28 months of age. The bands were detected with antibodies against indicated proteins. GAPDH was used as a loading control. Accumulations of HSP70, ubiquitin and p62 were prominent in the soleus muscle (arrows) and the diaphragm (arrowheads) of Dnajb4^F90L/+^ or Dnajb4^-/-^ mouse. GC, gastrocnemius; SOL, soleus; EDL, extensor digitorum longus; TA, tibialis anterior; HAM, hamstring; QF, quadriceps femoris; PS, paraspinal; DIA, diaphragm; PM, pectoralis major; TB, triceps brachii; TON, tongue; MAS, masseter

### Soleus is susceptible to Dnajb4 dysfunction

We reasoned that selective muscle involvement in patients may similarly occur in our mouse model. Therefore, to examine DNAJB4 expression and determine the susceptibility to DNAJB4 dysfunction in specific muscles in rodents, we performed western blotting using the lysate from 12 different muscles groups from wild-type, KI, and KO mice (Fig. 4d). This demonstrated that DNAJB4 was most highly expressed in the soleus muscles. This is consistent with published RNAseq data^18^ (http://muscledb.org). Furthermore, accumulations of HSP70, ubiquitin, and p62 were most prominent in the soleus muscles of Dnajb4^F90L/+^ or Dnajb4^-/-^ mouse, suggesting that the soleus requires higher levels of DNAJB4 to maintain proteostasis. This is consistent with the patient muscle pathology since the soleus muscle in mice contains a large number of type 1 fibers^19^. The subsequent analyses utilized the soleus muscle.

### Dnajb4^-/-^ mice develop muscle weakness, fibrosis, and p62-positive inclusions in the soleus muscles

We analyzed *ex vivo* muscle contraction of the soleus muscles of Dnajb4^-/-^ mice. Dnajb4^-/-^ mice had significantly lower peak isometric force and maximum tetanic force with or without cross sectional area (CSA)-normalization of the soleus muscles at the age of 28 months (Fig. 5a-h).

**Fig. 5.**
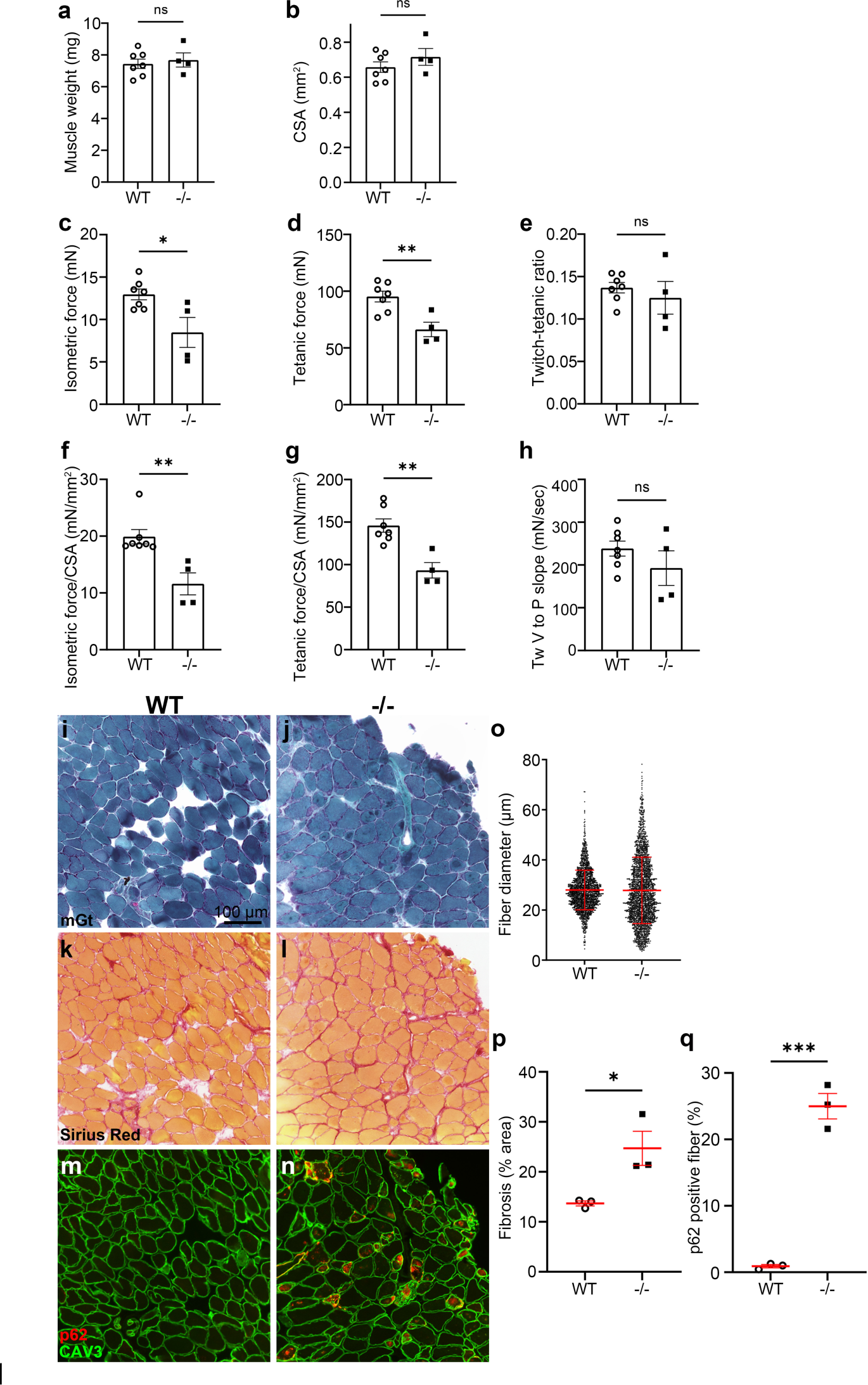
Assessment of *ex vivo* contraction and pathology of soleus muscles in Dnajb4^-/-^ mice at age of 28 months. **a**-**h** Muscle weight (**a**), cross sectional area (CSA) (**b**), and contractile forces (**c**-**h**) of the soleus muscles of Dnajb4^-/-^ (-/-) and Dnajb4^+/+^ (WT) at 28 months of age (n = 4-8 muscles from 2-4 mice per each group): (**c**) isometric twitch force, (**d**) maximum tetanic force, (**e**) the twitch-tetanic ratio between **c** and **d**, (**f**) specific isometric force normalized by CSA, (**g**) specific tetanic force normalized by CSA, and (**h**) the slope of valley (V) to peak (P) of isometric twitch force. **i**-**n** Muscle pathology and immunofluorescent images of the soleus muscles of Dnajb4^+/+^ (**i**, **k**, **m**) and Dnajb4^-/-^ (**j**, **l**, **n**) mice at age of 28 months. Muscle membranes are stained by anti-caveolin-3 (CAV3) antibodies (green). Scale bar = 100 μm (**i**-**n**). **o-q** Quantification of myopathological features of **i**-**n** (n = 4-8 muscles from 3-4 mice per each group): beeswarm plot of the fiber dimeter (**o**), area of fibrosis (**p**), ratio of p62 positive fibers (**q**). Values from each group are expressed as mean ± SEM except for fiber diameter (mean ± SD, **o**). A paired Student’s t-test for comparison between two groups was performed. *p < 0.05, **p < 0.01, ***p < 0.001, and ns, not significant. Scale bar = 100 μm (**i**-**n**). mGt, modified Gomori trichrome; Tw, twitch

Consistent with this weakness, mGt staining of the soleus sections from 28- month-old Dnajb4^-/-^ mice showed variation in fiber size, increased fibers with internal nuclei, and cytoplasmic inclusions compared to the age-matched wild type (Fig. 5i, j, o). On Sirius Red, the proliferation of interstitial connective tissues was seen (Fig. 5k, l, p). In addition, p62-positive fibers were increased (Fig. 5m, n, q)

### Dnajb4^F90L/+^ mice develop weakness, myofiber atrophy, fibrosis, and formation of p62-positive inclusions in the soleus muscles

To explore a DNAJB4 susceptible muscle, we performed *ex vivo* contraction using the soleus muscles from Dnajb4^F90L/+^. Both at 11 months and 28 months of age, absolute isometric force and CSA-normalized isometric force of the soleus were significantly lower in Dnajb4^F90L/+^ (Fig. 6a-h). Twitch-tetanic ratio and valley to peak slope of isometric force were lower in Dnajb4^F90L/+^, suggesting affected muscles slowly contracted on electric stimuli^20, 21^.

**Fig. 6.**
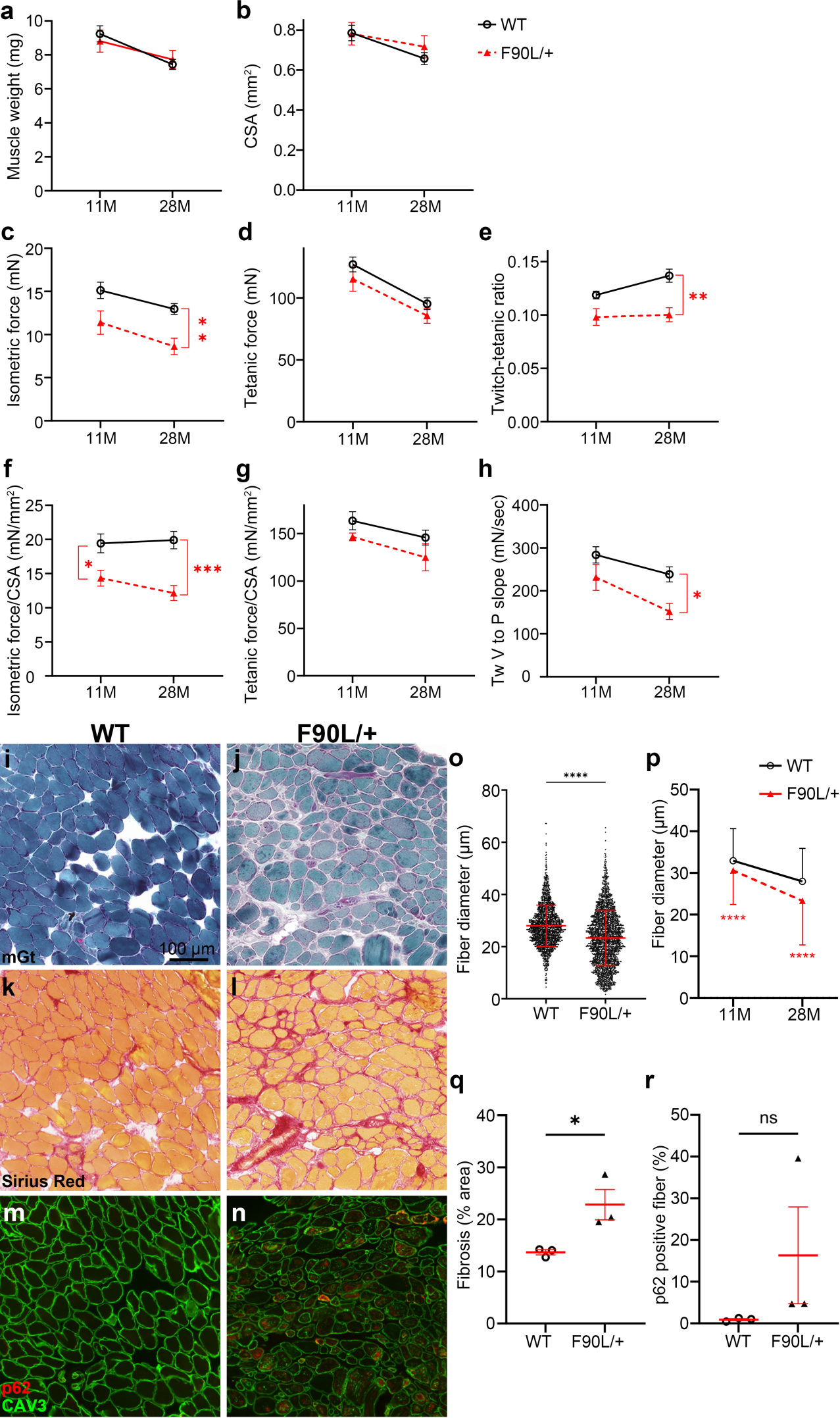
Assessment of *ex vivo* contraction and pathology of the soleus muscles in Dnajb4^F90/+^ mice at age of 11 and 28 months. Muscle weight. (**a**), cross sectional area (CSA) (**b**), and contractile forces (**c**-**h**) of the soleus muscles of Dnajb4^F90L/+^ (F90L) and Dnajb4^+/+^ (WT) at 11 months (11M) and 28 months (28M) of age (n = 7-8 muscles from 4 mice per each group). The data of Dnajb4^+/+^ at 28 months is identical to those in Fig. 5: (**c**) isometric twitch force, (**d**) maximum tetanic force, (**e**) the twitch-tetanic ratio between **c** and **d**, (**f**) specific isometric force normalized by CSA, (**g**) specific tetanic force normalized by CSA, (**h**) the slope of valley (V) to peak (P) of isometric twitch force. **i**-**n** Muscle pathology and immunofluorescent images of the soleus muscles of Dnajb4^+/+^ (**i**, **k**, **m**, identical images to Fig. 5i-n) and Dnajb4^F90L/+^ (**j**, **l**, **n**) mice at age of 28 months. Muscle membranes are stained by anti-caveolin-3 (CAV3) antibodies (green). Scale bar = 100 μm (**i**-**n**). **o**-**r** Quantification of myopathological features of **i**-**n** (n = 7-8 muscles from 4 mice per each group): fiber dimeter (**o**, beeswarm plot; **p**, time-course change), area of fibrosis (**q**), ratio of p62 positive fibers (**r**). Values from each group are expressed as mean ± SEM except for fiber diameter (mean ± SD, **o**, **p**). A paired Student’s t-test for comparison for comparison between two groups or a multiple testing comparison with multiple testing correction (Bonferonni) was performed. *p < 0.05, **p < 0.01, ***p < 0.001, ****p < 0.0001, and ns, not significant. mGt, modified Gomori trichrome; Tw, twitch

On muscle pathology of the soleus from 28-month-old Dnajb4^F90L/+^, we observed variation in fiber size, increased fibers with internal nuclei, and cytoplasmic inclusions compared to the age-matched wild type (Fig. 6j). Fiber size was significantly decreased at the age of 11 months, and the variation was greater at 28 months (Fig. 6o, p). We also found prominent fibrosis (Fig. 6l, q) and an increase in fibers with p62- positive inclusions (Fig. 6n, r) in the soleus of 28-month-old Dnajb4^F90L/+^ mice.

### Accumulation of Z-disc components and colocalization of HSP70

As mentioned above, we found the accumulation of proteins associated with the Z-disc, the molecular chaperone HSP70, and autophagy adaptor p62 in patient muscle. We analyzed the accumulation of these proteins in the muscles of Dnajb4^F90L/+^ and Dnajb4^-/-^ mice. In the soleus muscles of Dnajb4^F90L/+^ mice at 28 months, p62 was located in the center of the cytoplasmic inclusions observed on mGt, and was surrounded by DNAJB4, HSP70, and desmin (Fig. 7j). Interestingly, filamin C was present within the inclusion, but not colocalized with p62 (Fig. 7n). In the soleus muscles of Dnajb4^-/-^ mice at 28 months, as expected, immunoreactivity of DNAJB4 was significantly reduced (Fig. 7o, q). Otherwise, similar to the findings in Dnajb4^F90L/+^ mice, we observed desmin, filamin C, HSP70, and p62 at cytoplasmic inclusions (Fig. 7c, f, o-t).

**Fig. 7.**
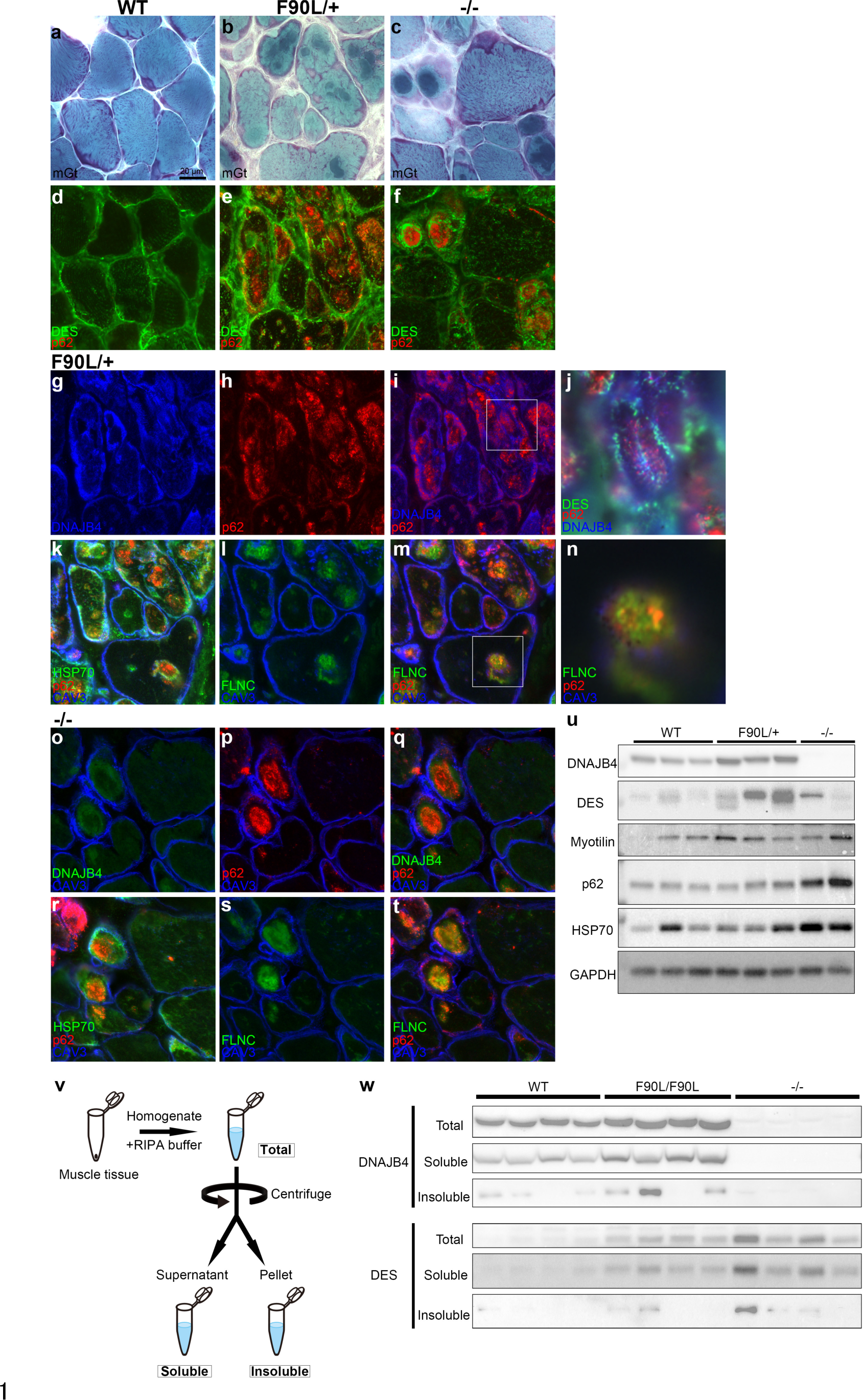
Protein accumulation in the soleus muscles from Dnajb4^F90L/+^ and Dnajb4^-/-^ mice at age of 28 months. **a**-**t** Modified Gomori trichrome (mGt) and immunofluorescent images staining for the indicated antibodies of the soleus muscles of Dnajb4^+/+^ (WT) (**a**, **d**), Dnajb4^F90L/+^ (F90L/+) (**b**, **e**, **g**-**n**) and Dnajb4^-/-^ (-/-) (**c**, **f**, **o**-**t**) mice at age of 28 months. **j**, **n** are magnified images of the white boxes of **i**, **m**, respectively. Muscle membranes are stained by anti-caveolin-3 (CAV3) antibodies (blue). Scale bar = 20 μm (**a-t**). **u** Immunoblot of lysates from the soleus muscles of Dnajb4^+/+^, Dnajb4^F90L/+^, and Dnajb4^-/-^ mice at 28 months of age. The bands were detected with antibodies against indicated proteins. GAPDH was used as a loading control. **v** Schematic diagram of the RIPA soluble/insoluble fractionation protocol. Gastrocnemius muscle lysates from Dnajb4^+/+^, Dnajb4^F90L/F90L^ (F90L/F90L), and Dnajb4^-/-^ mice at 13 months of age are separated into a total, soluble, and insoluble fraction. **w** Each fraction was immunoblotted for DNAJB4 and desmin. DES, desmin; FLNC, filamin C; HSP70, heat shock protein 70

To confirm the findings on immunohistochemistry, we immunoblotted using the lysate from the soleus muscle of Dnajb4^F90L/+^ mice at 28 months, and found increased levels of desmin, myotilin, and HSP70 (Fig. 7u). Notably, there was an increase in DNAJB4 protein levels. Muscles from Dnajb4^-/-^ had an increase in p62 (Fig. 7u).

Similar results were observed in the soleus muscles by immunohistochemistry and gastrocnemius muscles by immunoblot from younger Dnajb4^F90L/F90L^ or Dnajb4^-/-^ at the age of 13 months (Supplementary Fig. 4). Furthermore, we evaluated the solubility of proteins in the Dnajb4^F90L/F90L^ muscles by separating tissue lysates into soluble and insoluble fraction in RIPA buffer and found DNAJB4 was recovered in the insoluble pellet in muscles from Dnajb4^F90L/F90L^. Similarly, desmin in muscles both from Dnajb4^F90L/F90L^ and Dnajb4^-/-^ mice was also shifted to the insoluble form, suggesting they were aggregated (Fig. 7v, 7w).

### Ultrastructural observation revealed widespread myofibril disorganization

We further examined the myofibril structure in the soleus muscles of 13-month-old Dnajb4^F90L/F90L^ mice by electron microscopy. We found a wide range of cytoplasmic bodies along the longitudinal axis in myofibers, some spanning 40 sarcomeres, consistent with the larger area of p62 in the human patient muscle on immunostaining (Fig. 8a, b, d). The boundary between cytoplasmic bodies and surrounding myofibrils was distinct, but the electron dense materials located to the periphery were contiguous to the Z-disc of the normal-appearing myofibrils (Fig. 8a). Low-density cytoplasmic bodies contained amorphous material (Fig. 8d). Furthermore, under high magnification, degenerated thin and thick filaments were observed next to the intact Z-disc (Fig. 8c), suggesting the degeneration may begin at thin and thick filaments. Electron dense structures composed of wooly and sand-like pleomorphic structures were also observed (Fig. 8e).

**Fig. 8.**
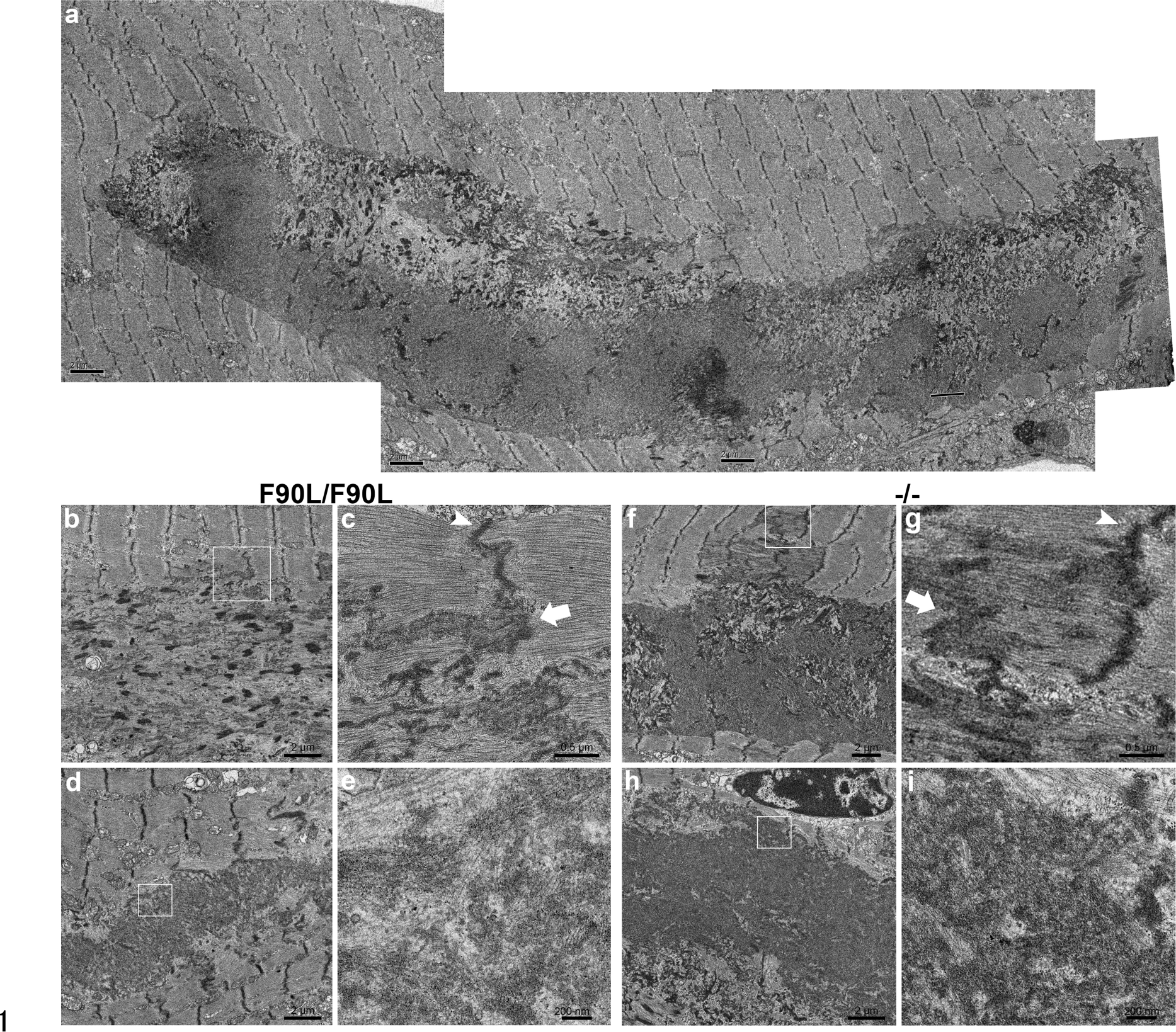
Ultrastructural findings in the soleus muscles from Dnajb4^F90L/F90L^ and Dnajb4^-/-^ mice at age of 13 months. Ultrastructural images of the soleus muscles from Dnajb4^F90L/F90L^ (**a**-**e**) and Dnajb4^-/-^ **(f**- **i). c**, **e**, **g**, **i** are magnified view of white boxes of **b**, **d**, **f**, **h**, respectively. **a** Combination of four separately taken pictures showing widespread cytoplasmic bodies along longitudinal axis in myofibers. **b**, **f**. Disorganized myofibrillar alignment with electron dense cytoplasmic bodies and the apparently clear border with surrounding myofibrils. **c**, **g** Degenerated thin and thick filaments (arrowhead) adjacent to the appearing-normal end of the Z-disc (arrowhead). **d**, **h** Low-density cytoplasmic bodies contained amorphous material. **e**, **i** Magnified images showing wooly and sand-like structures in the inclusions. Scale bar = 2 μm (**a**, **b**, **d**, **f**, **h**), 0.5 μm (**c**, **g**), 200 nm (**e**, **i**)

We also obtained similar ultrastructural findings in the soleus muscle of 28- month-old Dnajb4^-/-^ mice and found less dense inclusions surrounded by structures with electron dense materials similar to Z-disc (Fig. 8f, h), degenerated myofibrils next to normal Z-discs (Fig. 8g), and wooly sand-like materials within the magnified view (Fig. 8i).

### Diaphragms displayed cytoplasmic inclusion and myofibril destruction in Dnajb4^F90L/F90L^ and Dnajb4^-/-^ mice

Combining the type 1 fiber-predominant defects and the patients’ respiratory impairment led us to consider the possibility that the genetic alteration in DNAJB4 may affect the diaphragm. On mGt of the diaphragm from 13-month-old Dnajb4^F90L/F90L^ and Dnajb4^-/-^ mice, we observed a variation in fiber size and cytoplasmic inclusions compared to age-matched control mice (Supplementary Fig. 5a-c). On serial sections, NADH-TR showed that the fibers with cores corresponding to the cytoplasmic inclusions were found predominantly in type 1 fibers (Supplementary Fig. 5e,f).

Ultrastructural analysis using the diaphragm in 13-month-old Dnajb4^F90L/F90L^ and Dnajb4^-/-^ revealed cytoplasmic inclusions whose electron density were similar to those found in the soleus muscle of Dnajb4^F90L/F90L^ and Dnajb4^-/-^ mice (Supplementary Fig. 5g, i). Wooly sand-like materials in the magnified view were also observed in the diaphragms (Supplementary Fig. 5h,j).

### Mutant DNAJB4 showed toxic gain of function through the interactions with HSP70 in cultured cells

To evaluate the function of wild-type or DNAJB4-F90L against aggregate-prone proteins, we transfected both wild-type or mutant DNAJB4, and mutant polyglutamine (polyQ) or mutant desmin into C2C12 cells. The rates of cells with aggregates of polyQ and desmin were reduced by both wild-type and mutant DNAJB4, showing mutant DNAJB4 is effective against polyQ-derived aggregation or desmin accumulation (Fig. 9a-d).

**Fig. 9.**
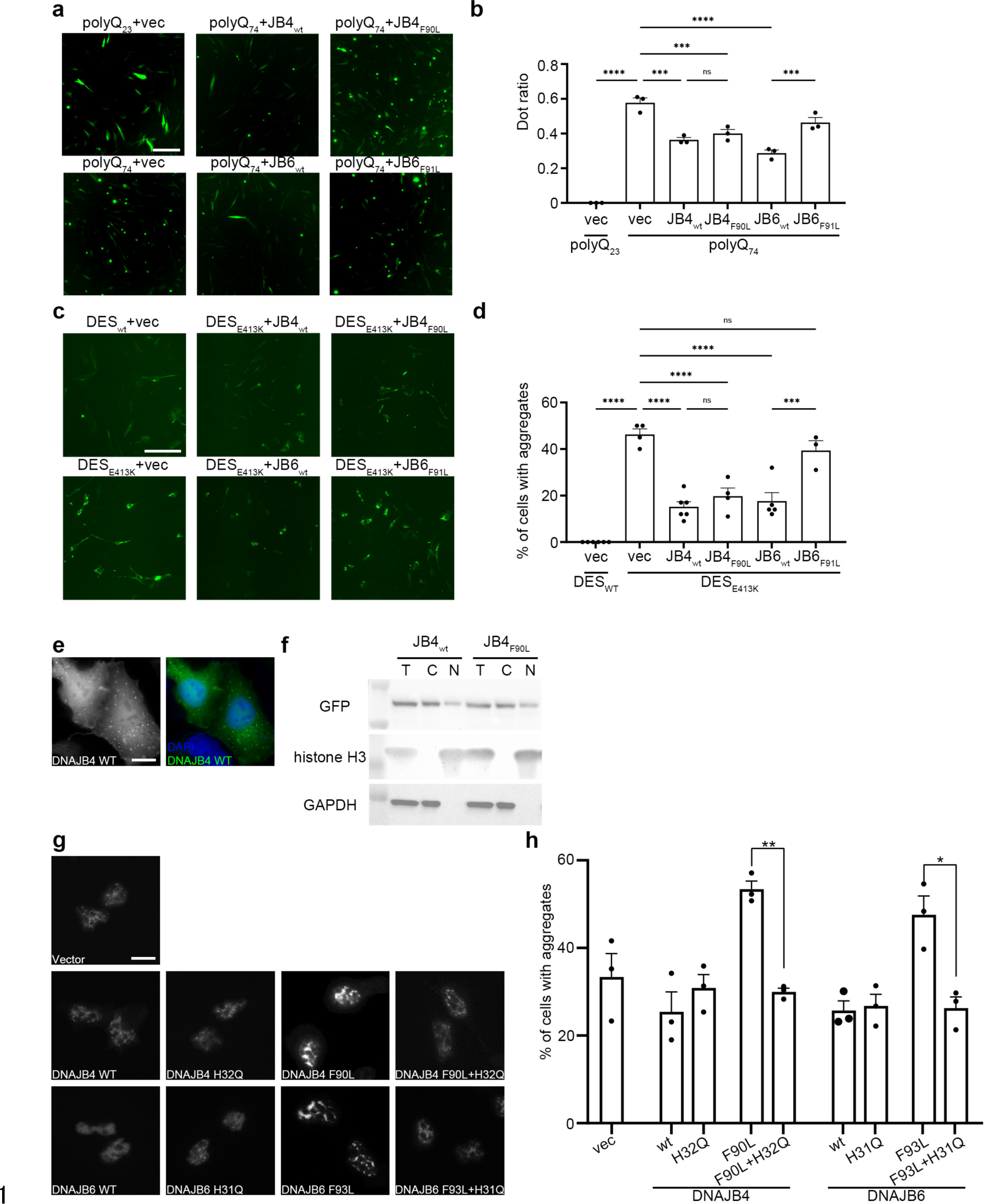
F90L mutation in *DNAJB4* impairs TDP-43 disaggregation, which is corrected with a second H32Q mutation. Representative fluorescence images of the GFP-polyQ. (**a**), GFP-desmin (**c**), and mCherry-TDP-43-positive nuclei (**g**). Scale bar = 200 μm (**a**, **c**), 10 μm (**g**). Quantification of the percentage of transfected cells with polyQ aggregates (**b**), mutant desmin (E413K) aggregates (**d**), and TDP-43 aggregates (**h**). **e** GFP-tagged wild-type (WT) DNAJB4 are localized in nucleus and cytoplasm in HeLa cells. Scale bar = 10 μm (**e**). **f** Immunoblot of total (T), cytoplasmic (C), and nuclear (N) fractions from HeLa cells. Blots were probed with anti-GFP, histone H3, or GAPDH antibodies. Anti-histone H3 and GAPDH antibodies were used to identify nuclear and cytoplasmic subcellular fractions, respectively. At least three independent experiments were performed in each assay. Values from each group are expressed as mean ± SEM. A multiple testing comparison with multiple testing correction (Tukey) was performed. *p < 0.05, **p < 0.01, ***p < 0.001, and ****p < 0.0001. DES, desmin; polyQ, polyglutamine

Because LGMDD1-mutant DNAJB6b showed a toxic gain-of-function effect through the interaction with HSP70 *in cellulo* models ^22^, we hypothesized that DNAJB4 mutant also have a similar toxic gain-of-function effect. To test this, we first analyzed the subcellular localization of wild-type and mutant DNAJB4 in HeLa cells and found that DNAJB4-F90L did not change its localization (Fig. 9e, f). Then, we co-transfected mCherry-tagged TDP-43 with plasmid constructs containing GFP-tagged wild-type or mutant DNAJB4 with or without an additional H32Q mutation that abolishes HSP70 association into HeLa cells. Although disease-causing F90L mutation increased the rate of cells with TDP-43 nuclear aggregates after heat shock, the escalation was abrogated by the additional H32Q mutation, suggesting F90L mutant DNAJB4 also has a toxic gain-of-function effect through its interaction with HSP70 similar to mutant DNAJB6 ^22^(Fig. 9g, h).

## Discussion

In this study, we characterize a chaperonopathy presenting as a late onset distal myopathy and protein aggregates in myofibers, associated with dominant mutations in the *DNAJB4*. Our KO mouse model showed myopathy, suggesting that DNAJB4 may be essential for skeletal muscle integrity. Moreover, our KI (F90L) mouse model reproduced the human disease phenotype. In addition, we gained new insight regarding muscle susceptibility to chaperone dysfunction.

Affected patients had a distinctive distal myopathy phenotype. The initial symptoms in the third or fifth decade of life are asymmetric hand grip weakness, especially thumb weakness with atrophy of thenar and hypothenar muscles. These are rarely observed in other chaperonopathies^6, 23^ or distal myopathies^24^. However, intriguingly, there is some overlap between distal myopathies caused by mutations in *DNAJB4* and *FLNC* in terms of atrophy of hand muscles and grip weakness^25^. In addition, muscle MRI revealed selective involvement of the rectus femoris and posterior thigh muscles and asymmetric fatty replacement, which is a pattern not previously described in other myopathies^26, 27^ and in contrast to DNAJB6-related myopathies in which rectus femoris is relatively spared^28, 29^. Thus, this muscle MRI pattern is highly unique and may be useful to distinguish DNAJB4 myopathy from other neuromuscular diseases, although further study including more patients is needed.

Furthermore, our results suggest that positional susceptibility within skeletal muscles may underlie selective muscle involvement in certain human myopathies. In patients with neuromuscular diseases, certain muscle groups are selectively involved, and others are relatively spared^30, 31^. Muscle imaging plays an essential role in identifying genetically different conditions, although little is known about the mechanism of the selective muscle involvement of neuromuscular diseases^27^. Here, among skeletal muscles of our model mice, the most affected muscles by the DNAJB4 dysfunction were the soleus and diaphragm, which have the highest percentage of type 1 myofibers^32, 33^. On patient muscle imaging, the more severely affected muscles were antigravity muscles, such as the biceps femoris, soleus, and paraspinal muscles, which have the highest percentage of type 1 fibers in humans^32^. In addition, the thenar and hypothenar muscles atrophied in the patients include the adductor pollicis brevis and short palmaris muscles, which are also known to be type 1 fiber predominant^32, 34, 35^.

Together, the selective susceptibility of the muscles to genetic alterations may contribute to selective muscle involvement. This susceptibility could be associated with type 1 fiber predominant involvement in DNAJB4 myopathy, but further studies are necessary to clarify the mechanism of selective muscle involvement.

DNAJB4 myopathology is also unique. In addition to fiber size variation and fibrosis, large cytoplasmic inclusions were observed in myofibers. Those are morphologically distinct from rimmed vacuoles only seen in small atrophic fibers.

Previously, we have characterized the muscle pathology of patients harboring F93L in *DNAJB6*, in which cytoplasmic inclusions and rimmed vacuoles were also observed^28^. Interestingly, the morphology, localization in myofibers, and accumulated protein profile of the inclusions were different between the two diseases. In DNAJB6 myopathy, inclusions are smaller, located subsarcolemmally, and co-stained with DNAJB6, myotilin, desmin, HSPB8, BAG3, Hsc70, and STUB1. In contrast, in DNAJB4 myopathy, inclusions are larger, located in the center of myofibers, and co-stained with DNAJB4, Hsp70, p62, filamin C, and desmin. Since DNAJB6 interacts with members of the chaperone-assisted selective autophagy complex (CASA)^28, 36^, it is suggested that the molecular pathology of DNAJB6 myopathy is mediated through impaired functions of CASA. However, in DNAJB4 myopathy, we failed to detect members of CASA and LC3 in the deposits and also failed to detect autophagic vacuoles under electron microscopy. Since the localization of DNAJB4 and DNAJB6 in normal muscles is quite similar, DNAJB4 should have a chance to interact with members of CASA. Further studies, especially proteomic identification of the accumulated proteins in deposits, are required to answer this question.

As the predominant expression of DNAJB4 is in striated muscles, patients with the variant in *DNAJB4* developed a pure myopathy. Skeletal muscle is an organ where protein synthesis is generally active, and many stresses including oxidative, metabolic and mechanical, continuously occur^37^. Molecular chaperones take part in protein quality control by chaperoning unstable proteins in skeletal muscles^3^, and their loss or dysfunction may be harmful, producing toxic misfolded proteins, which may become a seed for protein aggregation or propagate the misfolded conformation to other molecules by interaction with normally folded proteins^38–40^. In this study, we generated new mouse models for DNAJB4 myopathy. Our KO model showing myopathic phenotypes suggests that DNAJB4 is essential for muscle function and recessive mutations in *DNAJB4* could cause human myopathy in a loss of function manner.

Indeed, recessive mutations in *DNAJB4* have been identified in three families, demonstrating a myopathy with early respiratory failure (personal communication). Furthermore, our KI model mice recapitulated the muscle phenotypes of the human patients. The similar pathological features between the KI and KO mice imply that both the heterozygous variant and depletion of Dnajb4 cause similar effects on pathogenesis, suggesting F90L-DNAJB4 has a dominant negative effect on normal DNAJB4 to suppress the function. Additionally, the presence of larger cytoplasmic inclusions is characteristic of muscle pathology in human patients as well as model mice. Actually, those inclusions are p62-positive and extended for more than 30 sarcomeres in length in patient muscle, indicating the accumulated proteins are ubiquitinated and propagated along myofibers. Again, further studies on identification of ubiquitinated proteins in these cytoplasmic inclusions will clarify the functions of the HSP70-DNAJB4 chaperone system and the toxicity of misfolded proteins, which may be directly associated with the pathomechanism of this disease.

The *Dnajb4* gene-modified mice developed myopathy with age. This suggests that aging will be one of the key factors for developing this disease. Many molecules and pathways (protein synthesis, post translational modification, refolding, degradation) cooperatively work to maintain proteostasis in skeletal muscles, which are continuously exposed to stressful conditions^3^. Once one molecule/pathway is affected, other molecules/pathways can compensate^41, 42^, while with aging, these compensatory pathways gradually decline^43^. Whereas a defect in one pathway may be tolerated at a young age due to proteostatic compensation, at the older age, the declined compensatory system is no longer sufficient to maintain proteostasis resulting in protein aggregation^44, 45^.

The variant site, phenylalanine at 90, is within the conserved G/F domain in DNAJB4. Similar to the DNAJB6-F93L mutant, the DNAJB4-F90L mutant enhanced the formation of TDP-43-positive nuclear stress granules when expressed in cells, and the abrogation of its association with HSP70 suppressed TDP-43 stress granule formation^22^. These results suggested that the F90L variant in DNAJB4 has a gain-of- toxicity with HSP70 on TDP-43. On the other hand, interestingly, DNAJB4-F90L still retained comparable chaperone activities against mutant polyglutamine and desmin aggregates to the wild-type DNAJB4, suggesting that polyglutamine and desmin may be the client proteins of DNAJB4 and probably, these chaperone activities may be independent of the pathogenesis of the variant in cell experiments. We need further studies to elucidate the discrepancy between the results found *in cellulo* experiments and in animal models.

In conclusion, we discovered a form of chaperonopathies caused by a dominant variant in *DNAJB4*, which has distinctive clinical and myopathological features. This work also provides the basis of proteostasis specific to selective muscle groups.

## Methods

### Patients

Written informed consent was obtained from the affected or unaffected individuals. This study was approved by the institutional ethics committee of the National Center of Neurology and Psychiatry and Washington University in St. Louis. All the human materials used in this study were obtained for diagnostic purposes. The patients or their parents provided written informed consent for the use of the samples for research.

### Histological and immunohistochemical analyses for human and mouse samples

Histological analyses were performed as described previously ^46^. Isolated muscle samples were collected and then frozen in isopentane cooled in liquid nitrogen. Serial frozen sections (10-µm thick) were subjected to a battery of histochemical stains including hematoxylin and eosin, mGt, NADH-TR, or Sirius Red. For immunohistochemical analyses, tissue sections were fixed in acetone and blocked with 2% casein in phosphate-buffered saline. The primary antibodies used are as follows: anti-αB-Crystallin (Stressgen, 1:100), anti-alpha-actinin-2 (Sigma-Aldrich, 1:100), anti-caveolin-3 (Santa Cruz Biotechnology, 1:100), anti-desmin (abcam, 1:200), anti- DNAJB4 (Abgent, 1:100), anti-DNAJB4 (Proteintech, 1:100), anti-DNAJB6 (abcam, 1:100), anti-filamin C (Cloud-Clone, 1:100), anti-filamin C (Santa Cruz Biotechnology, 1:50), anti-HSPB8 (Abnova, 1:167), anti-HSC70 (abcam, 1:250), anti-HSP70 (abcam, 1:250), anti-p62 (PROGEN, 1:100), and anti-ubiquitinylated proteins (Enzo Life Sciences, 1:100). The sections were probed in appropriate secondary antibodies. Images were collected with a BZ-X-700 microscope (Keyence, Osaka, Japan). For single-fiber diameter, sections were probed with caveolin-3 (Santa Cruz Biotechnology, 1:100) followed by appropriate secondary antibodies. Images from whole areas of five or six sections of soleus muscles of three mice of each group were captured at ×200 magnification. From these images, individual fiber diameter was measured from 500- 700 fibers with J-image software (National Institutes of Health), taking note of the shortest diameter.

### Electron microscopy

Electron microscopic analysis was performed as previously described ^47^. The quadriceps femoris muscle of II-5 was fixed in 2.5% glutaraldehyde and post-fixed with 2% osmium tetroxide. Ultrathin sections stained with uranyl acetate and lead citrate were observed under a transmission electron microscope (FEI, Hillsboro, USA).

### Whole exome sequencing and Sanger sequencing

Genomic DNA was isolated from peripheral blood (II-1, II-5, II-7, III-4, III-6) or autopsied liver (II-3) using standard techniques. We performed whole exome sequencing for affected individuals (II-1, II-5, and II-7) as previously described^47–,49^(Supplementary Table 2).

Genetic variation was annotated with the NCNP in-house pipeline, and detection of known polymorphisms using gnomAD, dbSNP150, Human Genetic Variation Database, and Integrative Japanese Genome Variation Database for Japanese genetic variants. Candidates of variants identified in exome sequencing were evaluated in other affected and unaffected members by Sanger sequencing on an ABI Prism 3130 DNA Analyzer (Applied Biosystems Waltham, MA).

### Generation of the *Dnajb4* KI and KO lines

*Dnajb4* KI and KO mouse lines were generated via zygote electroporation utilizing the CRISPR/Cas9 system described previously^50, 51^. We designed the CRISPR RNA (crRNA) to target the coding sequences near c.270C > A (p.F90L) (Supplementary Fig. 2a). The crRNA and the trans-activating crRNA (tracrRNA) were chemically synthesized (FASMAC), and recombinant Cas9 protein (EnGen Cas9 NLS from *S. pyogenes*) was purchased from New England Biolabs. The single-stranded DNA (ssDNA) donor containing the c.270C > A substitution along with two silent mutations flanked by 50 nucleotides homology arms on each side was subsequently designed and chemically synthesized by Eurofins Genomics. The crRNA, tracrRNA, Cas9 protein, and ssDNA donor were electroporated into mouse one-cell-stage zygotes to obtain founder KI and KO mice, which were generated secondarily by the CRISPR/cas9 system. The founders were crossed with wild-type mice to obtain F1 generations. The

Dnajb4 KI (Dnajb4^F90L/+^) and Dnajb4 KO (Dnajb4^+/-^) mice were backcrossed on the C57BL/6J strain. Homozygous Dnajb4 KI (Dnajb4^F90L/F90L^) mice were generated by crossing Dnajb4^F90L/+^ mice. Homozygous Dnajb4 KO (Dnajb4^-/-^) mice were generated by crossing heterozygous Dnajb4^+/-^ mice. The mice genotypes were identified by the Sanger sequence (Supplementary Fig. 2b). Data only from female mice were used in this study. Animals were kept in a barrier-free, specific pathogen-free grade facility on a 12-hour light/12-hour dark cycle and had free access to standard chow and water. All animal study procedures in this study were approved (Approval No. 2022014, R4-03) by and carried out within the Ethical Review Committee on the Care and Use of Rodents in the National Institute for Neuroscience, National Center of Neurology and Psychiatry.

### Grip tests

Grip tests were performed as previously described^52^. The grip strength of combined forelimbs/hindlimbs was measured by ten consecutive trials using an MK-380S grip strength meter (Muromachi Kikai Co., Ltd) for each mouse. The resulting measurement was recorded, and the average taken by excluding the highest and lowest values was determined to give the strength score. For every time point and strain, at least four animals were used.

### Muscle contractile properties analysis

We analyzed the *ex vivo* muscle contraction of the soleus, gastrocnemius, and tibialis anterior muscles as previously described ^52, 53^.

### Single fiber isolation

For immunofluorescence, the soleus muscles from wild-type C57BL/6J mice were dissected and fixed as previously described^54^. Briefly, after fixation with 4% paraformaldehyde for 30 minutes at room temperature, single fibers were obtained by manual teasing.

### Second harmonic generation

We performed second harmonic generation imaging as previously described^55^. Briefly, we electroporated 25 μg of constructs expressing GFP-tagged DNAJB4-WT or –F90L into FDB muscles of C57BL/6J mice after anesthesia. Mice were euthanized one week after electroporation, and whole ankles were dissected and fixed with 4% paraformaldehyde. The mouse feet were placed onto a customized viewing pane to facilitate imaging with a Zeiss LSM 880 Airyscan two-photon microscope. A single excitation wavelength of 800 nm was used for 2-photon microscopy and recorded using a ×40 1.2 NA water immersion objective.

### Western blot

Frozen muscle tissues were sliced and solubilized in the SDS-PAGE buffer containing 125 mM Tris, 2% sodium dodecyl sulfate (SDS), 10% glycerol, 5% 2-mercaptoethanol, pH 6.8, or homogenized using RIPA buffer (50 mM Tris-HCl pH8.0, 150mM NaCl, 1% NP-40, 0.5% sodium deoxycholate, 0.1% SDS) supplemented with protease inhibitor cocktail (Roche or MilliporeSigma, MO). SDS-polyacrylamide gel electrophoresis was conducted following Laemmli’s method^56^. Lysates were centrifuged at 16,000 *g* for 10 minutes. Protein concentrations were measured using the Bio-Rad Protein Assay Kit or BCA Protein Assay Kit (Thermo Fisher Scientific). Equal amounts (25 μg or 10 μg) of protein were separated on 4-20% SuperSep Ace (FUJIFILM Wako Pure Chemical Corporation, Osaka, Japan), polyacrylamide gradient gel (Bio-Rad), or 12% SDS-PAGE gels, and transferred onto polyvinylidene difluoride or nitrocellulose membranes. The membrane was incubated with the following primary antibodies: anti-αB-Crystallin (Enzo Life sciences, 1:1000), anti-alpha Tubulin (1:800), anti-BAG3 (abcam, 1:500), anti-desmin (Dako, 1:500), anti-DNAJB4 (Abgent, 1:1000), anti-DNAJB4 (Proteintech, 1:1000), anti-GAPDH (Cell Signaling Technology, 1:1000), anti- HSP70/HSP72 (Enzo Life sciences, 1:1000), anti-p62 (Proteintech, 1:1000), anti-p62 (Novus Biologicals, 1:1000), and anti-TDP-43 (Proteintech, 1:1000). The membranes were washed, probed in appropriate HRP-labeled secondary antibodies, washed, and then made to react with Immobilon Western Chemiluminescent HRP Substrate (MilliporeSigma). Detection was done by ImageQuant LAS4000 (GE Healthcare) or G:Box Chemi XT4, Genesys, version 1.1.2.0 (Syngene). The signal intensity was quantified with ImageJ software (NIH).

### Solubility assay

Soluble and insoluble fractions were analyzed as previously reported^57, 58^. Fresh gastrocnemius muscles were collected and frozen in liquid nitrogen-cooled isopentane. Tissue was homogenized in RIPA buffer (50 mM Tris-HCl pH8.0, 150mM NaCl, 1% NP-40, 0.5% sodium deoxycholate, 0.1% SDS) and protease inhibitor cocktail (MilliporeSigma). Homogenates were sonicated and centrifuged at 13,000 *g* for 10 minutes at 4°C. Protein concentrations of the supernatant were measured using BCA Protein Assay Kit, and all samples were diluted to the same concentration for fractionation. An aliquot of the supernatant was collected and labeled the total fraction. The additional supernatant was centrifuged at 100,000 *g* for 30 minutes at 4°C, and this supernatant was collected and named the soluble fraction. The pellet was washed once with 200 μL of RIPA buffer, resonicated, and recentrifuged. Then the supernatant was removed and labeled the insoluble fraction. The insoluble fraction was dissolved in 50 μL of urea buffer (7M urea, 2M thiourea, 4% CHAPS, 30mM Tris pH 8.5) and resonicated. Each sample was analyzed by Western blotting for each fraction.

### Construction of DNAJB4 and DNAJB6 mutants and transfection into HeLa or C2C12 cells

For expression analysis, the open reading frame of human DNAJB4, DNAJB6, and desmin were inserted into expression vectors pcDNA6 V5-His A (Invitrogen) using BamHI/XhoI. Expanded poly-Q (polyQ23 and polyQ74) in human huntingtin were digested with HindIII/EcoRI and ligated into a vector pEGFP-C1 containing a GFP tag. For heat shock experiments, human DNAJB4 (Addgene) and DNAJB6 (Addgene) constructs were made as previously described^55^. The DNAJB4 and DNAJB6 cDNAs were digested with HindIII/XhoI and ligated into the vector pcDNA3.1 containing a GFP tag or a V5 tag. All mutations were introduced with the QuikChange Mutagenesis Kit (Agilent Technologies). Information regarding primer sequences, cloning conditions, and polymerase chain reaction is available upon request. We merged Human TDP43 to pCherry as previously described^59^.

### Nuclear/cytoplasmic fractionation

Nuclear/cytoplasmic fractionation was performed as previously described^60^. HeLa cells were harvested in trypsin/EDTA (MilliporeSigma) and washed twice with ice-cold PBS. An aliquot was collected, incubated in RIPA buffer, and labeled the total fraction. The additional cells were incubated in mild cell lysis buffer [20 mM Tris, pH 7.4, 10 mM KCl, 3 mM MgCl2, 0.1% NP-40, 10% Glycerol, protease inhibitor cocktail (Roche), and PMSF] for 10 min on ice and centrifuged at 2000 *g* for 10 min at 4 °C, and this supernatant was collected and named the cytoplasmic fraction. The pellet was resuspended with the same amount of mild cell lysis buffer and named the nuclear fraction. After adding 4× SDS-PAGE buffer to the total fraction, the cytoplasmic fraction, and the nuclear fraction, equal volumes of each fraction were analyzed by Western blotting with the following antibodies: anti-GAPDH (Cell Signaling Technology, 1:1000), anti-GFP (Cell Signaling Technology, 1:1000), anti-histone H3 (abcam, 1:1000).

### Cell culture, transfection, and heat shock in HeLa cells

We performed heat shock experiments as previously described ^55^. HeLa cells were grown in Dulbecco’s modified Eagle’s medium (DMEM, Gibco) supplemented with 10% fetal bovine serum, penicillin/streptomycin. HeLa cells were co-transfected with 1) GFP-DNAJB4 or GFP-DNAJB6 and 2) mCherry-TDP-43 constructs and maintained at 37°C for 24 hours. Cells were subjected to heat shock (42°C, 5% CO2) for 1 hour, then allowed to recover at 37°C for 2 hours.

### Cell culture, transfection, and immunohistochemistry in C2C12 myoblasts

C2C12 cells were grown in DMEM (Gibco) supplemented with 10% fetal bovine serum, penicillin/streptomycin. To evaluate the rate of cells with aggregate with DNAJB4 and DNAJB6, we co-transfected 1) DNAJB4 or DNAJB6 constructs and 2) GFP-tagged polyQ or desmin constructs into C2C12 cells using Lipofectamine LTX and Plus reagent (Thermo Fisher Scientific). The rate of the cells with aggregate was observed three days post-transfection.

### Statistics

All values were expressed as means ± SEM except for values of myofiber diameters (mean ± SD). The data were then subjected to a paired Student’s t-test for comparison between two groups, Dunnett’s, Tukey’s, or Bonferroni’s multiple comparison test for more than two groups as appropriate. Kaplan-Meier survival curves and the log rank test were used to compare cumulative survival. All analyses were performed with GraphPad Prism 9 (GraphPad Software). P value less than 0.05 was considered statistically significant.

### Data availability

The source data underlying Figs. 3, 4, 5, 6, 7, and 9 are provided in the Source Data file. The reported novel variant has been submitted Leiden Open Variation Database (LOVD, https://databases.lovd.nl/shared/variants, Variant ID: 0000871428). Any additional data that support the findings of this study are available from the corresponding author upon reasonable request.

## Acknowledgements

We thank the patients and their families for their cooperation. We also thank Hisayoshi Nakamura, Keiko Hiraki-Kamon, and Keiko Ishikawa for technical assistance. This study was supported by the Intramural Research Grant for Neurological and Psychiatric Disorders of the National Center of Neurology and Psychiatry under grant numbers 2-5 (S.H., I.N.), 2-6 (S.N.), 3-8 (M.T.), and 3-9 (S.N.); KAKENHI (19K17021) from the Japan Society for the Promotion of Science (M.I.); and Japan Agency for Medical Research and Development grant numbers 22ek0109490h0003 (A.I., S.H., S.N., I.N.).

## Author contribution

M.I., S.N., and I.N. designed and oversaw this study. M.I., S.N., Y.U.I., A.I., M.O., R.B., S.K.P., and T.I. performed experiments. M.I., S.N., A.I., and C.C.W. analyzed data. M.I., K.W., Y.H., T.S., M.T., Y.O., Y. T., and H.M. provided samples and clinical data. M.I., S.N., and C.C.W. wrote the manuscript. All authors contributed to and approved the final manuscript.

