## Supplementary Material for "Distinctive chaperonopathy in skeletal muscle associated with the dominant variant in *DNAJB4*"

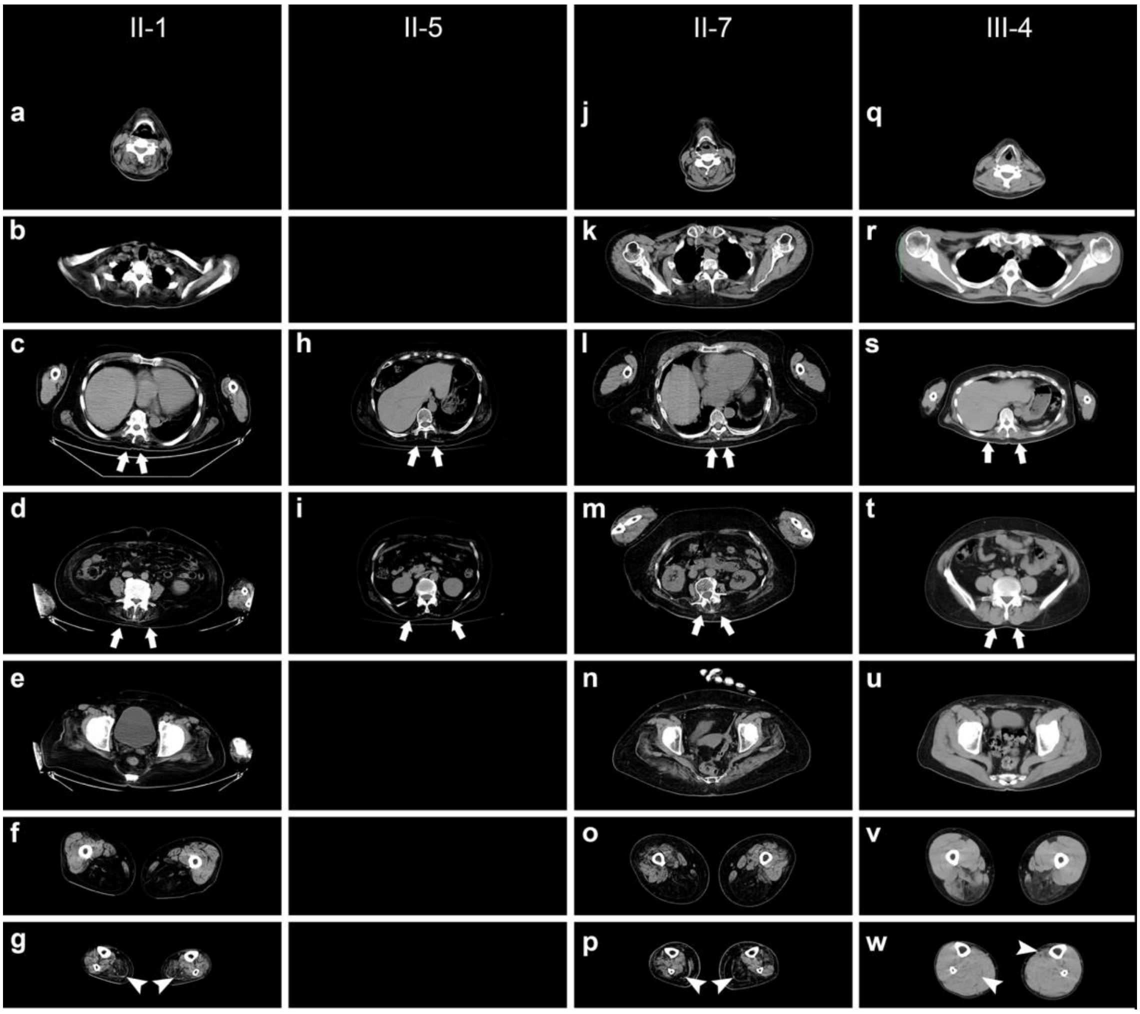


**Supplementary Fig. 1 Muscle CT images of the patients.**

CT images for II-1 (**a**-**g**) , II-5 (**h**, **i**), II-7 (**j**-**p**), and III-4 (**q**-**w**) at the age of 71, 68, 74, and 49, respectively: neck (**a,** **j**, and **q**), shoulder (**b**, **k**, and **r**), chest (**c**, **h**, **l**, and **s**), abdomen (**d**, **i**, **m**, and **t**), hip (**e**, **n**, and **u**), thigh (**f**, **o**, and **v**), and lower leg (**g**, **p**, and **w**). Arrows indicate the paraspinal muscles. Arrowheads indicate the soleus muscles

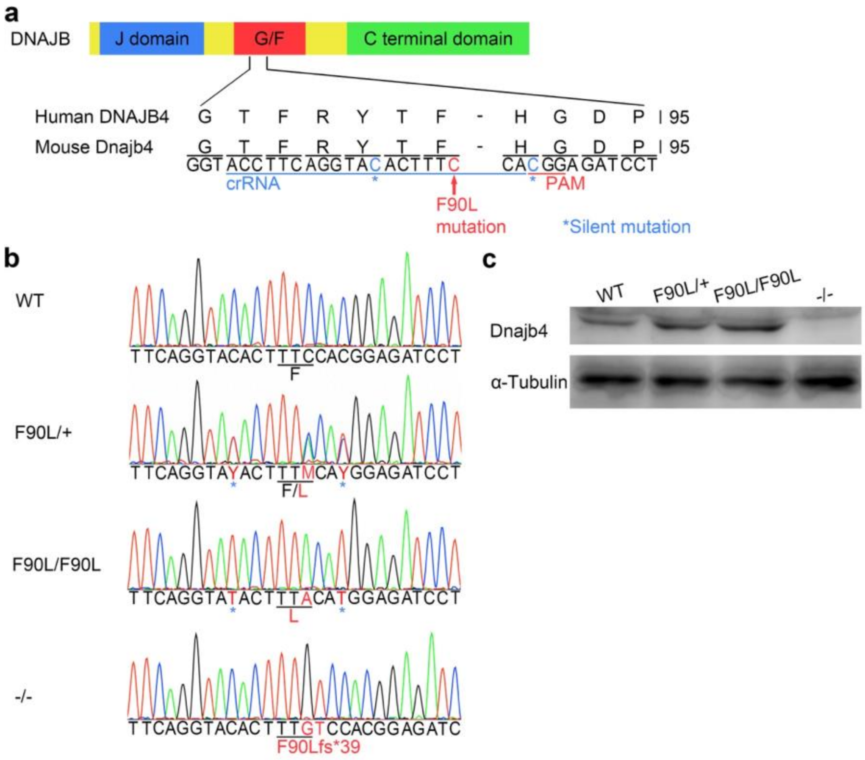


**Supplementary Fig. 2 Generation and verification of the Dnajb4$^{F90L/+}$, Dnajb4$^{F90L/F90L}$, and Dnajb4$^{-/-}$ mice.**

**a** CRISPR RNA (crRNA) was designed to target the coding sequences near c.270C>A (p.F90L) corresponding to human c.270T>A (p.F90L) variant in the glycine/phenylalanine-rich (G/F) domain of DNAJB4. Two silent mutations (blue) were additionally introduced to interrupt the protospacer adjacent motif (PAM) and crRNA sequences, preventing unwanted re-editing of the repaired template. **b** Electropherograms of each genotype verified the sequences of the F90L KI with silent mutations and the KO by frameshift at the *Dnajb4* locus. **c** Western blotting of proteins extracted from the quadriceps femoris muscles. The bands were detected with antibodies against DNAJB4. α-Tubulin was used as a loading control. WT, Dnajb4$^{+/+}$; F90L/+, Dnajb4$^{F90L/+}$; F90L/F90L, Dnajb4$^{F90L/F90L}$; -/-, Dnajb4$^{-/-}$

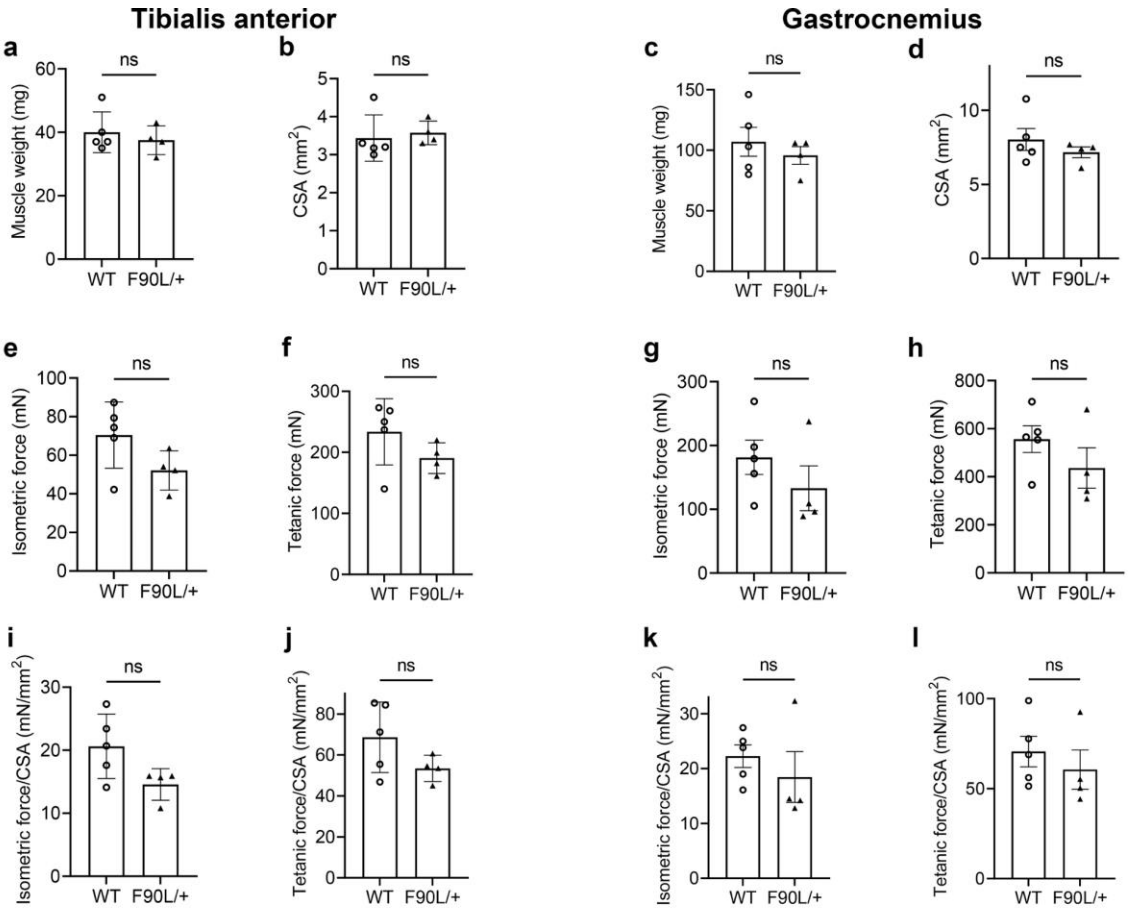


**Supplementary Fig. 3 Assessment of physiological parameters in Dnajb4$^{F90L/+}$ and Dnajb4$^{+/+}$ mice at age of 28 months.**

Muscle weight (**a**, **c**), cross sectional area (CSA) (**b**, **d**), and contractile forces (**e**-**l**) of the tibialis anterior muscles (**a**, **b**, **e**, **f**, **i**, **j**) and the gastrocnemius (**c**, **d**, **g**, **h**, **k**, **l**) of Dnajb4$^{F90L/+}$ (F90L/+) and Dnajb4$^{+/+}$ (WT) mice at 28 months of age (n = 4-5 per each group): (**e**, **g**) isometric twitch force, (**f**, **h**) maximum tetanic force, (**i**, **k**) specific isometric force normalized by CSA, (**j**, **l**) specific tetanic force normalized by CSA.

Values from each group are expressed as mean ± SEM. A paired Student's t-test for comparison between two groups was performed

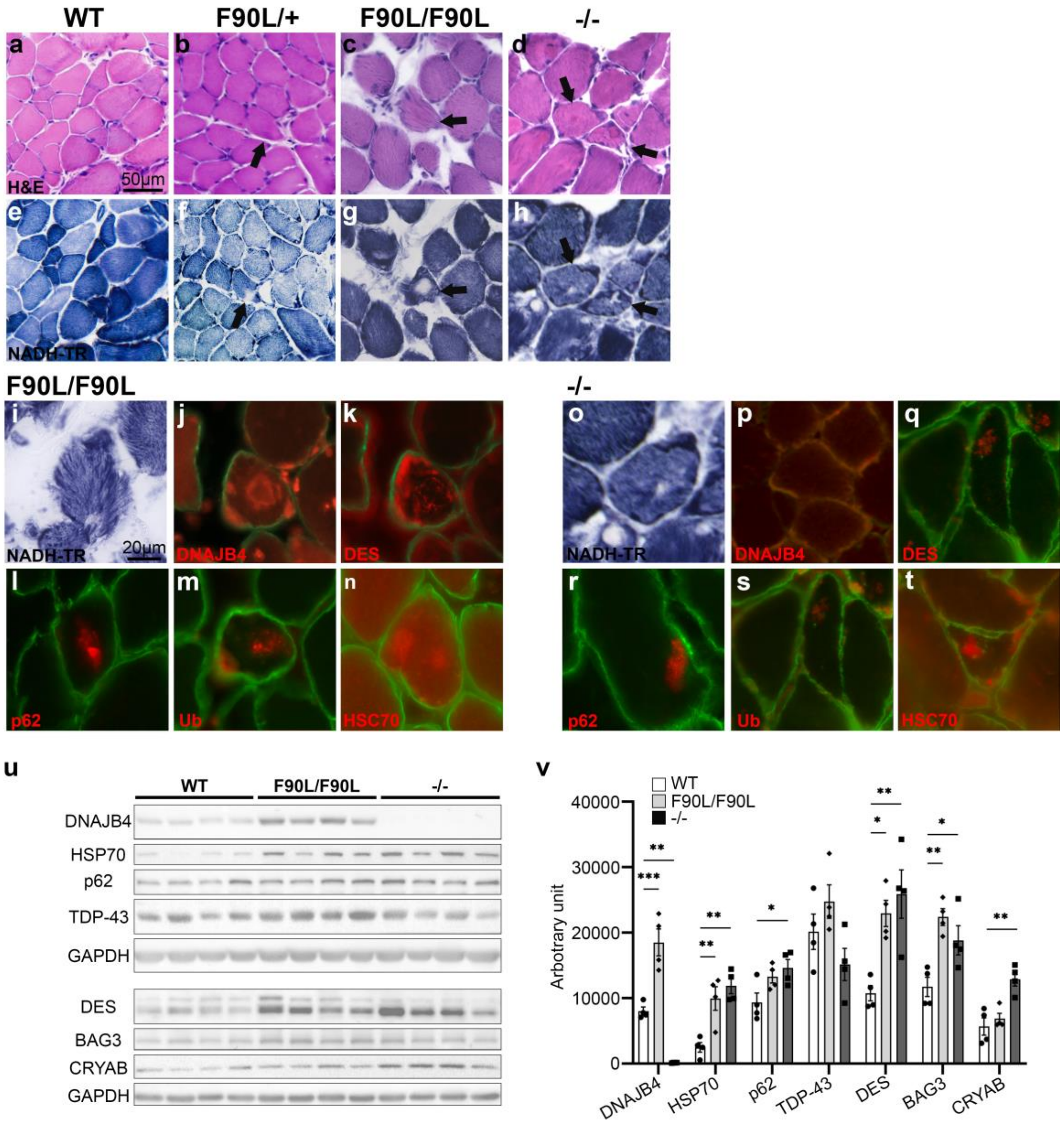


**Supplementary Fig. 4 Muscle pathology and protein accumulation in Dnajb4$^{F90L/+}$, Dnajb4$^{F90L/F90L}$, and Dnajb4$^{-/-}$ mice at the age of 13 months.**

Muscle pathology of the soleus muscles of Dnajb4$^{+/+}$ (WT) (**a**, **e**), Dnajb4$^{F90L/+}$ (F90L/+) (**b**, **f**), Dnajb4$^{F90L/F90L}$ (F90L/F90L) (**c**, **g**), and Dnajb4$^{-/-}$ (-/-) (**d**, **h**) mice at the age of 13 months. **a-d** Hematoxylin and eosin (H&E) staining. **e-h** Nicotinamide adenine dinucleotide dehydrogenase-tetrazolium reductase (NADH-TR) staining which are serial sections of **a-d**, respectively. H&E staining in Dnajb4$^{F90L/F90L}$ and Dnajb4$^{-/-}$show cytoplasmic inclusions (arrows) corresponding to the core-like structures in the serial section of NADH-TR (arrows) while H&E of Dnajb4$^{F90L/+}$ does not show inclusion in the corresponding fiber (arrow) showing core-like structure in NADH-TR. **i-t** NADH-TR and immunofluorescence staining for the indicated antibodies of the soleus muscles from Dnajb4$^{F90L/F90L}$ (**i-n)** and Dnajb4$^{-/-}$ (**o-t**) mice at the age of 13 months. Muscle membranes are stained by anti-caveolin-3 antibodies (green). **u** Immunoblot of lysates from the gastrocnemius muscles from four different Dnajb4$^{+/+}$, Dnajb4$^{F90L/F90L}$, and Dnajb4$^{-/-}$ mice at 13 months of age. The bands were detected with antibodies against indicated proteins. GAPDH was used as a loading control. **v** Densitometric quantification of four replicates of the blot is shown. Values from each group are expressed as mean ± SEM. A multiple testing comparison with multiple testing correction (Dunnett) was performed. *p < 0.05, **p < 0.01, and ***p < 0.001. BAG3, BAG cochaperone 3; CRYAB, alpha B crystallin; DES, desmin; FLNC, filamin C;

HSC70, heat shock cognate protein 70 ; HSP70, heat shock protein 70; TDP-43, TAR DNA binding protein; Ub, ubiquitin

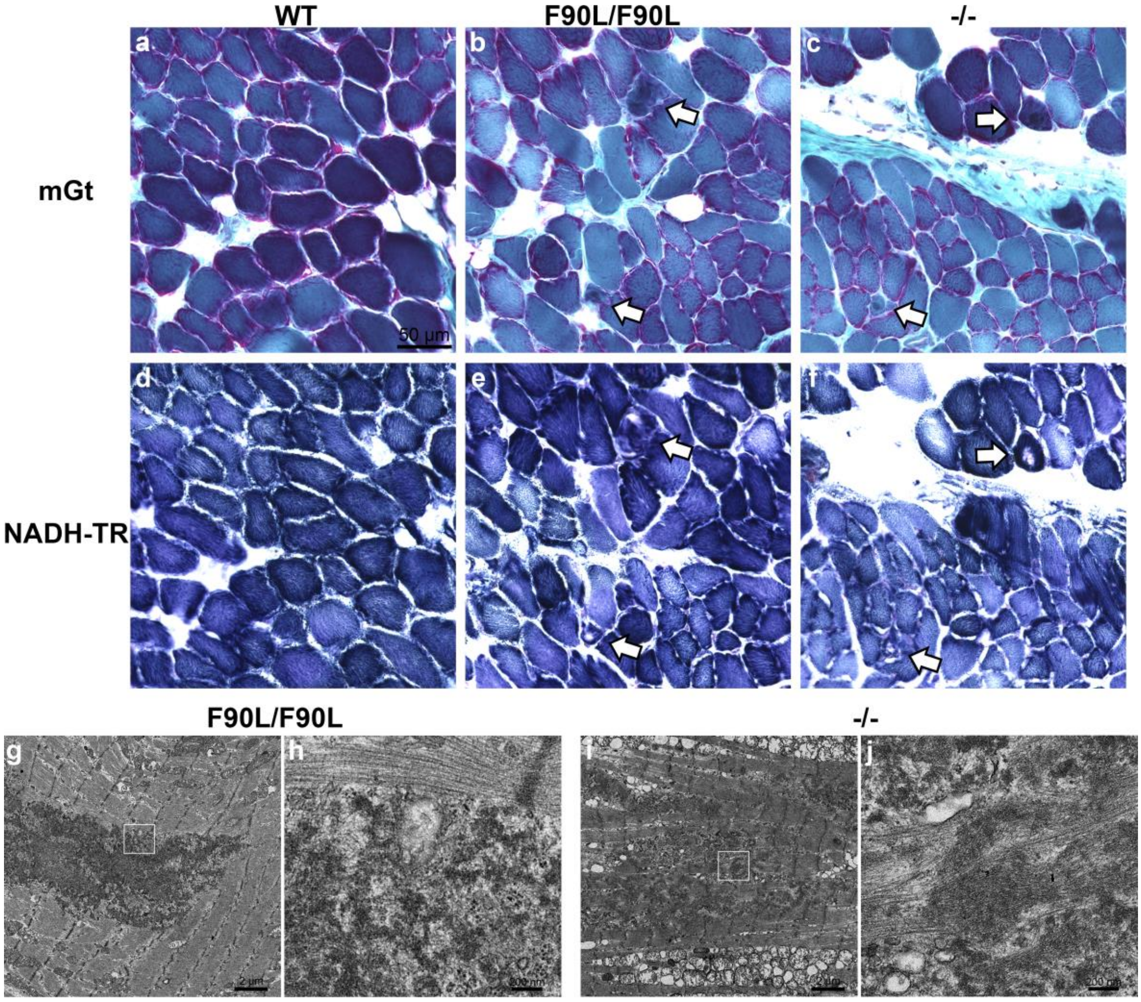


**Supplementary Fig. 5 Myopathological and ultrastructural features of diaphragms in Dnajb4$^{F90L/F90L}$ and Dnajb4$^{-/-}$ mice.**

Muscle pathology (**a**-**f**) and ultrastructural images (**g**-**i**) of diaphragms from Dnajb4$^{+/+}$ (WT) (**a**, **d**) Dnajb4$^{F90L/F90L}$ (F90L/F90L) (**b**, **e**, **g**, **h**) and Dnajb4$^{-/-}$ (-/-) (**c**, **f**, **i**, **j**) mice at 13 months of age. **b**, **c** Modified Gomori trichrome (mGt) showing cytoplasmic inclusions (arrows). **b** Nicotinamide adenine dinucleotide dehydrogenase-tetrazolium reductase (NADH-TR) stainings are serial sections of **a**-**c**, showing core-like structures (arrows). **g**, **i** Electron dense cytoplasmic bodies next to the apparently normal Z-discs. **h**, **j** Magnified views of white boxes in **g**, **i** showing wooly and sand-like structures in the inclusions. Scale bar = 50 μm (**a-f**), 2 μm (**g**, **i**), 200 nm (**h**, **j**)

### Supplementary Table 1. Genotype and clinical features of the family members

| | I-1 | II-1 | II-3 | II-5 | II-7 | III-4 | III-6 |
|---|---|---|---|---|---|---|---|
| **Genotype** | NA | c.270T>A (heterozygous) | NA | c.270T>A (heterozygous) | c.270T>A (heterozygous) | c.270T>A (heterozygous) | Ref/ref |
| **Sex** | Male | Male | Male | Female | Female | Male | Female |
| **Age of Onset** | late 20s | around 20 | 40s hand grip weakness | late 30s | Around 40 | Around 40 | No symptoms |
| **Age at last exam** | 71 (death) | 71 (death) | NA | 75 (death) | 74 | 49 | 45 |
| **Phenotype description** | distal myopathy | distal myopathy | distal myopathy | distal myopathy | distal myopathy | distal myopathy | |
| **Symptoms onset** | weakness of thumbs | hand grip weakness, ROM restriction of thumbs | hand grip weakness | decreased grip strength | hand grip weakness | hand grip weakness | |
| **Cause of death (age)** | prostate cancer (71) | respiratory failure at age (71) | NA | aspiration pneumonia and respiratory failure (75) | | | |
| **History of disease (Age—symptoms)** | NA | late 20s—asymmetric weakness of thumbs, hand grip weakness, weakness of limbs (from distal to proximal)<br>late 30s—difficulty in climbing upstairs<br>40s—difficulty in standing up from chairs, lordosis<br>50s—difficulty in raising upper limbs<br>62—bedridden, respiratory insufficiency | 40s—hand grip weakness<br>50s—difficulty in standing up and walking | late 30s—weakness of thumbs, hand grip weakness<br>40s— limb weakness from distal to proximal muscles<br>60s—using wheel chair | around 40—asymmetric hand grip weakness<br>44—equinus foot, Achilles tendon lengthening, limb weakness from distal to proximal muscles<br>68—shortness of breath on exertion<br>74—home oxygen therapy | around 25—much more difficulty in running than before<br>around 40, asymmetric hand grip weakness<br>47—difficulty in standing up without hand support | No symptoms |
| **Gait** | NA | | abnormal | 60s— non-ambulatory | 68—non-ambulatory without a walker | Normal | Normal |
| **Motor milestones** | Normal | Normal | Normal | Normal | Normal | Normal | Normal |
| **Hand grip weakness** | Yes | Yes | Yes | Yes | Yes | Yes | No |
| **Atrophy of thenar and hypothenar muscles** | Yes | Yes | Yes | Yes | Yes | Yes<br>(especially in adductor pollicis brevis) | No |
| **Contracture of fingers (flexed position)** | NA | Yes | NA | Yes | Yes | No | No |
| **Scoliosis** | NA | NA | NA | Yes | Yes | No | No |
| **Facial /Ophthalmological involvement** | No | No | No | No | No | No | No |
| **Cognition** | Normal | Normal | Normal | Normal | Normal | Normal | Normal |
| **Respiratory involvement** | No | Yes | No | Yes | Yes | No | No |
| **Cardiac involvement** | No | No | No | No | No | No | No |
| **Other** | prostate cancer | | | breast cancer<br>Chronic lower back pain | Chronic lower back pain | Chronic lower back pain | |
| **CK (Max)** | NA | 669 | NA | 866 | 646 | 1285 | NA |
| **EMG** | NA | NA | NA | myogenic | NA | myogenic | NA |
| **NCS** | NA | Normal | NA | Normal | NA | Normal | NA |

NA, not available; Ref, reference sequence; CK, creatine kinase; EMG, electromyography; NCS, nerve conduction study

**Supplementary Table 2. Summary of Exome sequencing performance and variant analysis**

| | II-1 | II-3 | II-4 |
|---|---|---|---|
| Number of alignment reads | 28,944,957 | 99,562,848 | 30,202,710 |
| Mean coverage[a] | 64.1268 | 162.64 | 68.2739 |
| % bases above 10x[b] | 90.79 | 96.41 | 97.19 |
| Filter passed SNVs[c] | 77394 | 86156 | 82090 |
| Filter passed indels[c] | 17457 | 16090 | 16720 |
| Exonic + Splicing | 22545 + 12331 | 23676 + 13271 | 23353 + 12818 |
| Absent in public databases[d] | 851 | 902 | 889 |
| Compatible with AD inheritance | 17 | | |
| Co-segregated among all family members examined | 2 | | |

[a] The mean depth of mapped reads

[b] Percentage of bases covered with more than 10 reads

[c] Variants that passed GATK filtering

[d] Absent or extremely low (< 0.000001) in gnomAD, dbSNP150, Human Genetic Variation Database, and Integrative Japanese Genome Variation Database
